# Movie Watching for Individual Functional Brain Network Estimation in Adolescents

**DOI:** 10.64898/2026.10.08.757407

**Authors:** Kate J. Godfrey, Gracie J. Grimsrud, Julian S.B. Ramirez, Allison Corlett, Kerry Houlihan, Nora Byington, Martin Gell, Sarah Kretsch, Nico U.F. Dosenbach, Deanna M. Barch, Evan M. Gordon, Deanna J. Greene, Bryon A. Mueller, Richard F. Betzel, Damien A. Fair, Christine A. Conelea, Brenden Tervo-Clemmens, Steven M. Nelson

**Author notes:** Denotes equal contribution (Christine A. Conelea, Brenden Tervo-Clemmens, Steven M. Nelson).

## Abstract

Precision functional mapping (PFM) reveals stable individual differences in brain functional network organization and holds promise for advancing neuropsychiatric clinical care. PFM requires sufficient high-quality and densely sampled functional MRI data at the individual level that is typically acquired during the so-called resting state while passively fixating on a crosshair. However, crosshair fixation acquisitions can be challenging especially for pediatric or patient populations, where alternatives like movie watching might be used to improve data quality and reduce participant burden. Here, we compare the use of resting state crosshair fixation and movie watching fMRI data to derive individual-specific functional brain network topography using PFM data (28–38 minutes of low-motion fMRI data per-participant per-condition) in nineteen adolescents (12-18 years) with a range of transdiagnostic psychiatric diagnoses. Network assignments were largely conserved between crosshair fixation and movie watching, with a smaller subset of reproducible network reassignments which were overwhelmingly border shifts between neighboring networks. This suggests largely conserved global structure of functional network topography between crosshair fixation and movie watching, though border shifts can compound into variable estimates of cortical network size particularly for movie positive (language, salience, and parietal memory) and movie negative networks (default mode, action mode). Functional network assignments derived from movies and crosshair were also similarly individual-specific and consistent within common neuromodulation targets including dorsolateral prefrontal cortex and somatomotor cortex. Key results were replicated in independent pediatric consortia data (Healthy Brain Network, *n* = 466, 6-22 years) and reproduced across independent PFM data sessions. Acknowledging individual differences and level of analysis, we conclude that movies are a viable alternative to resting state crosshair fixation for individual network definitions.

## INTRODUCTION

Functional MRI (fMRI) has shown that the human brain is organized into large-scale functional networks, supporting coordinated cognitive, affective, sensory, and motor processes (Biswal et al., 1995; Smith et al., 2009; Power et al., 2011; Yeo et al., 2011; Petersen & Sporns, 2015; Betzel & Bassett, 2017). Functional networks have historically been defined at the group-level by averaging across individuals; however, group averaging obscures individual features of functional networks that are considered critical for understanding individual differences in behavior (Bijsterbosch et al., 2018; Keller et al., 2023; Ladwig et al., 2026) and informing neuromodulatory interventions (Fox et al., 2013; Cash et al., 2021; Cole et al., 2022; Cash & Zalesky, 2024). To overcome these limitations, recent work has emphasized the development of precision functional mapping (PFM) protocols which leverage advancements in fMRI acquisition, denoising, and scan duration to facilitate high-reliability measurement of brain function in single individuals (Laumann et al., 2015; Gordon, Laumann, Gilmore, et al., 2017; Gratton et al., 2018; Lynch et al., 2020; Moser et al., 2025). Prior PFM studies demonstrate that functional brain networks in individuals reflect a mixture of common underlying spatial and functional properties of the group and trait-like individual features (Gordon, Laumann, Adeyemo, & Petersen, 2017; Gratton et al., 2018; Lynch et al., 2026), and that individual features of functional networks relate to individual differences in cognition and behavior (Bijsterbosch et al., 2018; Keller et al., 2023; Lynch et al., 2024; Vaghi et al., 2025; Ladwig et al., 2026).

Identification of individual functional brain networks with PFM holds particular promise for advancing ‘precision psychiatry’ through individually informed brain-based diagnosis and intervention (Gratton et al., 2020; Labonte et al., 2024; Demeter & Greene, 2025). PFM in adults has been used to identify individual differences in functional network topography (vertex-wise network assignments) that associate with psychiatric symptoms, including reports of salience network expansion in adults with major depressive disorder (Lynch et al., 2024) and obsessive-compulsive disorder (Vaghi et al., 2025). The use of PFM to define topography-based individual functional networks can also advance clinical care with brain network guided neuromodulation where stimulation is delivered to an individually defined optimal target site (c.f., (Fox et al., 2013; Cash et al., 2021; Lynch et al., 2022; Ren et al., 2026)). This is particularly true in dorsolateral prefrontal cortex, which is a common non-invasive neuromodulation target across psychiatric conditions (Li et al., 2022; Kan et al., 2023) and exhibits varied organization across individuals (Mueller et al., 2013; Fox et al., 2013; Ladwig et al., 2026). Importantly, studies considering PFM for precision psychiatry have largely focused on adults despite adolescence representing a critical window for emergent psychopathology (Paus et al., 2008; Uhlhaas et al., 2023), the increased use of neuromodulation in pediatric-focused clinical care and trials (El-Shahawy et al., 2025; C. A. Conelea & Lieske, 2026; Srivastava et al., 2026), and suggestions that heightened neuroplasticity during adolescence could be leveraged for neuromodulation treatments (C. A. Conelea & Lieske, 2026). It is therefore essential to consider whether the same individual principles of functional networks described in adult clinical cohorts are observed in adolescent clinical cohorts to support clinical applicability of PFM in earlier developmental stages.

Applications of PFM in developmental samples remain exceedingly sparse. One potential contributing factor is that PFM is typically applied using so-called resting state acquisitions, collected while passively fixating on a crosshair, which can be challenging for developmental cohorts (Vanderwal et al., 2019). Movie watching is one alternative which likely reduces participant burden and can decrease pediatric head motion (Vanderwal et al., 2015; Greene et al., 2018; Frew et al., 2022) therefore improving the quantity and quality of low-motion functional data (Van Dijk et al., 2012; Power et al., 2012; Satterthwaite et al., 2012) which is an essential consideration for PFM given fMRI reliability increases as a function of scan duration (Laumann et al., 2015; Gordon, Laumann, Gilmore, et al., 2017; Noble et al., 2017; Demeter et al., 2025; Rai et al., 2025; Ooi et al., 2025). Initial studies have demonstrated that individual functional networks in childhood and early adolescence can be independently derived using either resting state crosshair fixation (Demeter et al., 2025) or naturalistic stimuli including a movie and non-narrative clips (Rai et al., 2026) providing evidence towards the use of movies for individual functional network topography definitions.

Despite increased interest in movie watching for improved tolerability and feasibility of PFM, the effects of movie watching compared to standard crosshair fixation on individual functional network topography specifically (i.e., vertex-wise functional network assignments) have not been demonstrated in a pediatric clinical cohort. This is a critical consideration for use of movies in clinical contexts given one core goal of PFM is individual topography-based neuromodulation (c.f., (Fox et al., 2013; Cash et al., 2021; Lynch et al., 2022; Ren et al., 2026)). While prior work in adults comparing crosshair fixation and task regressed functional data suggest relatively minimal influences of acquisition condition on functional network topography (Kraus et al., 2021; Du et al., 2025) it’s unclear whether similarly minimal condition effects would be observed for movie watching. Movies in comparison to task stimuli are highly complex, irregular, and dynamic across high-level narrative features and low-level visual and auditory stimulus variations (Vanderwal et al., 2019); and movies evoke distributed alterations across sensory, cognitive control, attention, and association networks (Bottenhorn et al., 2018). However, despite these and other reported movie effects on functional network dynamics (for reviews, see (Vanderwal et al., 2019; Sonkusare et al., 2019; Jääskeläinen et al., 2021)) whether these dynamic changes between movie watching and crosshair fixation also manifest as variations in individual functional network topography remains unknown. Establishing whether individualized functional network topography is preserved across movie watching and crosshair fixation is critical for the long-term translational and basic-science goals of PFM, which depend on robust and generalizable representations of individual network organization. Toward these long-term clinical and basic-science goals, we examined movie watching and crosshair fixation as distinct acquisition states, rather than treating movie watching as a task, given the complexity of movie stimuli and the importance of identifying individualized network topography that generalizes across acquisition contexts.

Here we aim to advance PFM in developmental cohorts by evaluating individual functional brain network topography derived from fMRI data collected during passive movie watching and crosshair fixation acquisitions. To mirror the potential real world applicability of PFM in developmental and clinical cohorts, we collected a novel PFM dataset by recruiting adolescents with diagnoses for which neuromodulation treatment is available (major depressive disorder, (Srivastava et al., 2026)) or being tested in ongoing research (tic disorders, generalized anxiety disorder, obsessive compulsive disorder, autism spectrum disorder, and attention deficit hyperactivity disorder; (El-Shahawy et al., 2025)). Within this group we collected precision and high-reliability fMRI data while adolescents passively viewed movies (40 total minutes) and crosshair (40 total minutes) across two independent data collection sessions (80 total minutes of functional data per-participant). We demonstrate that whole cortex functional networks derived from movie watching and crosshair fixation acquisitions are similarly individual-specific and test-retest reliable, suggesting both conditions can be used for individual functional network topography estimation. We then describe common and specific features of functional network topography across viewing conditions, identifying that the majority of the cortical surface has conserved vertex-wise network assignments between movies and crosshair with a smaller set of reproducible network reassignments that are predominantly boundary changes between neighboring networks. We further identify that functional brain network topography is predominantly driven by condition-invariant group organization and trait-like individual differences, rather than differences between movies and crosshair fixation, though with notable regional variation across clinical neuromodulation targets. We conclude by demonstrating key results from our novel PFM dataset can be replicated in an independent pediatric consortia dataset (Healthy Brain Network). This work supports movie watching for PFM in certain contexts, while establishing key considerations for individual variability, regional differences, and condition-dependent effects in certain types of analyses.

## RESULTS

We collected a novel precision functional imaging dataset by recruiting youth with neurodevelopmental and psychiatric disorders that have prior indication for neuromodulation interventions that could be enhanced with PFM. The Mapping Individual Networks Transdiagnostically (MINT) (*n*=19, 12-18 years) neuroimaging data analyzed here includes 80 total minutes of functional magnetic resonance imaging (fMRI) for each individual collected across two independent data sessions for resting state crosshair fixation (40 total minutes; white crosshair on a black background) and movie watching (40 total minutes; concatenated *Pixar* short movies for children). Key study results were replicated using independent consortia data from the Healthy Brain Network (HBN) (*n*=466, 6-22 years) which included 10 minutes of fMRI for resting state crosshair fixation (white crosshair on a black background) and 13 minutes of fMRI across two movies (*Despicable Me* and *The Present*) collected during a single data session (Alexander et al., 2017). In alignment with the MINT precision data, the HBN consortia data included participants with neurodevelopmental and psychiatric diagnoses. Detailed demographics and sample characteristics for participants included in neuroimaging analyses are fully summarized in the supplemental materials for the MINT precision (Table S1) and HBN consortia data (Table S2).

### Functional data quality and quantity across movie watching and crosshair fixation

We first examined whether movie watching improved functional data quality by reducing head motion or increasing post-censored functional data quantity, which are essential considerations for the reliable identification of functional networks in individuals (Laumann et al., 2015; Gordon, Laumann, Gilmore, et al., 2017). When considering the MINT precision data prior to participant exclusions for fMRI data quality (*n*=28 total) movie watching significantly reduced head motion (mean framewise displacement in mm; *d* = -0.35, *p* < 0.04) while marginally improving data retention (post-censored minutes at FD < 0.2mm; *d* = 0.28, *p* = 0.08) (Figure 1, Table S3). In the HBN consortia data prior to fMRI data quality exclusions (*n*=1280), with a larger sample across a broader age range, movie watching significantly reduced head motion (mean framewise displacement; *d* = -0.19, *p* < 0.001) (Figure 1, Table S3). Examination of data quality outcomes by age in the HBN data suggests that benefits of movie watching to reduce head motion were primarily observed in younger participants which replicates prior reports (Greene et al., 2018; Frew et al., 2022) (Figure S1). These findings support prior work emphasizing the benefits of movie watching on data quality, particularly for younger pediatric groups.

**Figure 1.**
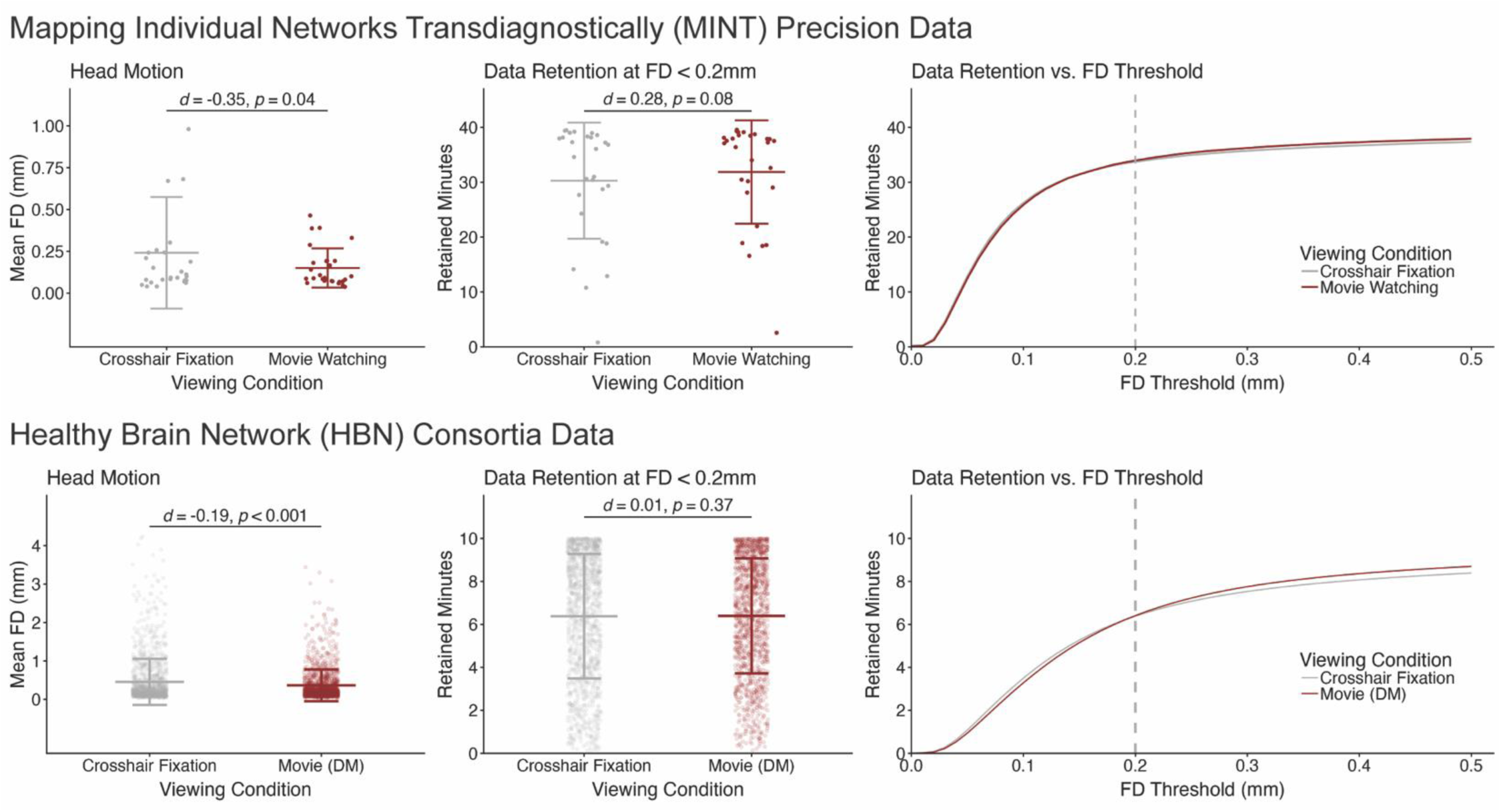
Functional data quality and quantity across movie watching and resting state crosshair fixation in precision and consortia data. Movie watching significantly reduced head motion (mean framewise displacement in mm, meanFD) in both the MINT precision (meanFD; crosshair mean = 0.24, movie mean = 0.15) and HBN consortia data (meanFD; crosshair mean = 0.45, movie mean = 0.36). Movie watching had a minimal, non-significant, effect on data retention at the study threshold of FD < 0.2mm (post-censored minutes) in both the MINT precision (post-censored minutes; crosshair mean = 30.28, movie mean = 31.85) and HBN consortia data (post-censored minutes; crosshair mean = 6.38, movie mean = 6.40). Descriptive and inferential statistics for condition effects on functional data quality and quantity outcomes are fully summarized in the supplemental materials (Table S3). FD: framewise displacement; Movie (DM): Movie (Despicable Me).

In our remaining neuroimaging analyses we aimed to increase reliability and control for motion effects, while also allowing for reproducibility analyses across independent data sessions, by including participants from the MINT precision data with 28-38 total minutes of low-motion functional data for each viewing condition and at least 10 minutes of low-motion functional data per condition at each independent data session. We included participants from the HBN consortia data if they had at least 8 minutes of low-motion functional data for each viewing condition. In the neuroimaging analysis sub-samples (MINT precision *n*=19, HBN consortia *n* = 466) viewing condition effects on functional data quality outcomes were not significant (all *p* > 0.2) (Figure S2).

### Functional Networks Derived from Movies or Crosshair are Individual-Specific and Reliable

We first examined the MINT data to compare crosshair fixation and movie watching for deriving individualized and reliable functional networks. Individual functional network topography was derived for cortical vertices using the template matching network detection algorithm (Gordon, Laumann, Adeyemo, Gilmore, et al., 2017). Template matching takes a dense connectivity matrix as input and calculates the spatial overlap of all vertex seedmaps to each template network; which produces an intermediate vertex-to-network similarity matrix (59k cortical vertices by 15 template networks); and derives final network topography using a winner-take-all heuristic where vertices are assigned to the network with the maximum similarity (for more details, see methods and Figure S3). Functional brain networks derived from both crosshair fixation and movie watching were examined for individual specificity (greater same individual over different individual similarity, on vs. off diagonal in Figure 2A) and test-retest reliability (same individual similarity across independent data sessions).

**Figure 2.**
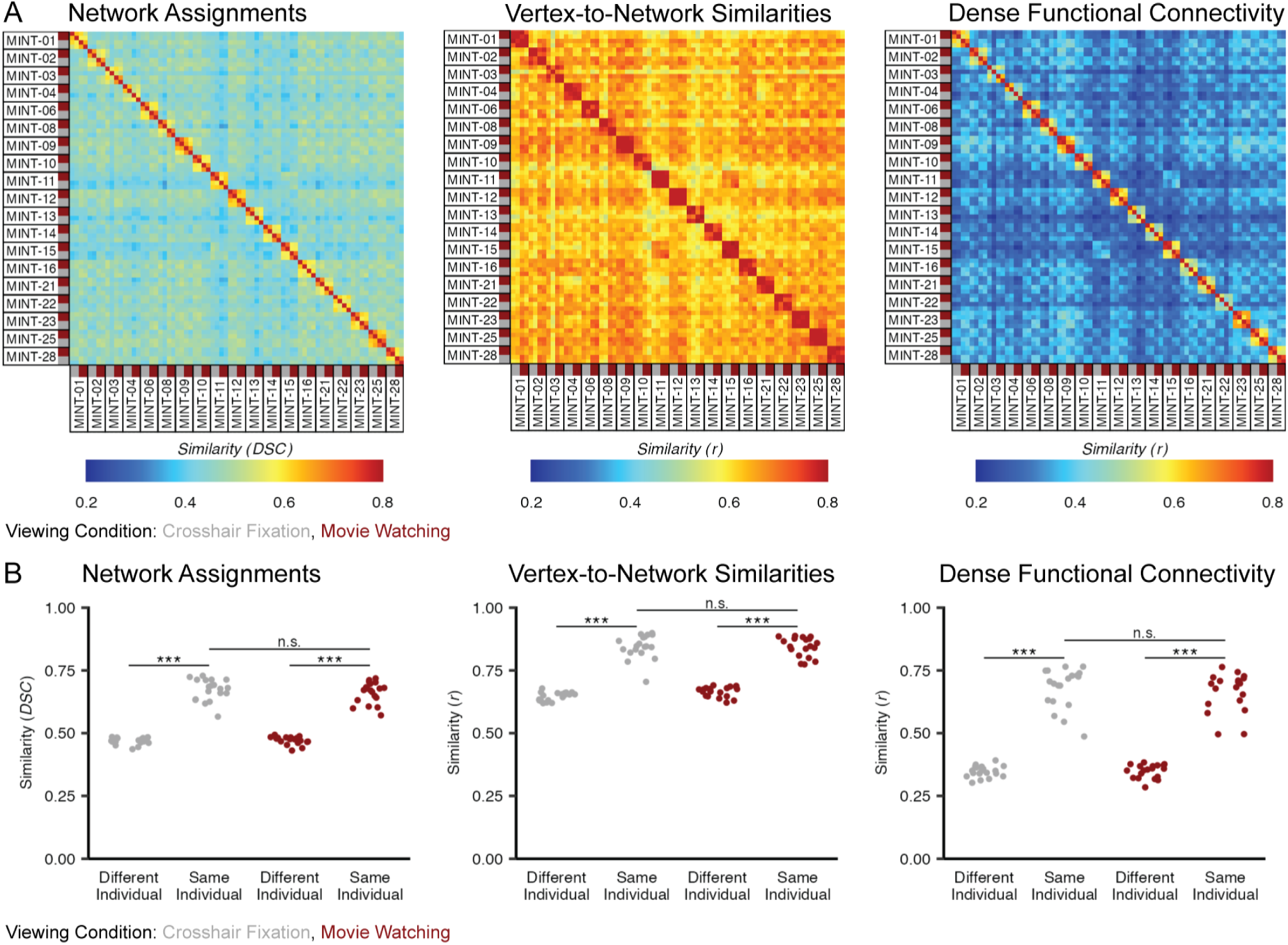
Large-scale functional networks derived with movie watching and crosshair fixation are similarly individual-specific and test-retest reliable. A) Similarity across all possible observation pairs in the MINT precision data (2 viewing conditions x 2 sessions x 19 participants) were derived for network assignments (network assignment vectors, Dice coefficient), template matching intermediate vertex-to-network similarity matrices (vectorized upper triangle, linear bivariate correlation), and dense functional connectivity matrices (vectorized upper triangle, linear bivariate correlation). B) The similarity of functional brain networks from the same individual (test-retest similarity across independent data sessions) were greater than the mean similarity of each individual to all different individuals across network assignments, vertex-to-network similarity matrices, and dense functional connectivity (all *p* ≤ 0.001). Functional brain networks derived from movie watching and crosshair fixation additionally did not differ in test-retest reliability across independent data sessions for network assignments, vertex-to-network similarity matrices, or dense functional connectivity (mean *p* > 0.24).

Functional brain networks in the MINT precision data derived from crosshair fixation or movie watching were individual-specific, such that repeat observations from the same individual were more similar than the mean similarity to different individuals. This improved same individual over different individual similarity was observed for the primary outcome of functional network topography (crosshair fixation: *t*(18) = 20.00, *p* < 0.001; movie watching: *t*(18) = 22.13, *p* < 0.001); the template matching intermediate vertex-to-network similarity matrices (crosshair fixation: *t*(18) = 19.20, *p* < 0.001; movie watching: *t*(18) = 22.56, *p* < 0.001) and dense functional connectivity (crosshair fixation: *t*(18) = 19.19, *p* < 0.001; movie watching: *t*(18) = 21.64, *p* < 0.001) (Figure 2B). This suggests that either movie watching or crosshair fixation can be used to identify individual differences to an equivalent magnitude. To test this, we further derived a per-participant relative individualization measure for crosshair fixation and movie watching (i.e., within each condition the per-participant test-retest same individual similarity minus the mean similarity to all other individuals) which did not differ in movie watching compared to crosshair fixation for the primary outcome of functional network topography (*t*(18) = -1.89, *p* = 0.07) or dense functional connectivity (*t*(18) = -1.57 *p* = 0.13) though relative individualization for template matching intermediate vertex-to-network similarity matrices was slightly higher for crosshair fixation (*t*(18) = -2.35 *p* = 0.03).

Functional brain networks in the MINT precision data derived from crosshair fixation and movie watching additionally did not differ in test-retest reliability across independent data sessions (mean = 6 days apart) for functional network topography (*t*(18) = 1.22, *p* = 0.23), template matching intermediate vertex-to-network similarity matrices (*t*(18) = 0.03, *p* = 0.98), or dense functional connectivity (*t*(18) = 1.25, *p* = 0.22) (Figure 2B). See also spatial distribution of reliable network assignments by condition at individual vertices (Figure S4).

### Movie Effects on Network Topography Are Driven by Shifting Boundaries at Network Borders

We next used the MINT precision data to visualize the consistency and variability of functional network topography (vertex-wise winner-take-all network assignments) between crosshair fixation and movie watching. When considering absolute magnitude of all possible consistent and discrepant functional network assignments across the entire cortex, approximately two thirds of the cortical surface (% of total cortical surface in mm^2^) had conserved network assignments across conditions (mean = 62.80, range = 57.79—63.73) while approximately one third of the cortical surface was discrepant (mean = 37.20, range = 32.37—42.21) (Figure 3). With a strict reproducibility criterion of identifying the same vertex-wise reassignment at each independent testing session, taking advantage of the MINT test-retest design, approximately 10% of the cortical surface was reproducibly reassigned (% of total cortical surface in mm^2^; mean = 8.95, range = 5.10–14.01) (Figure S4). This suggests that while a proportion of reassignments may be attributable to measurement error, a subset are highly reproducible across data sessions. Though we note that vertices with reliable network assignments during crosshair fixation and movie watching do not perfectly overlap (Figure S5).

**Figure 3.**
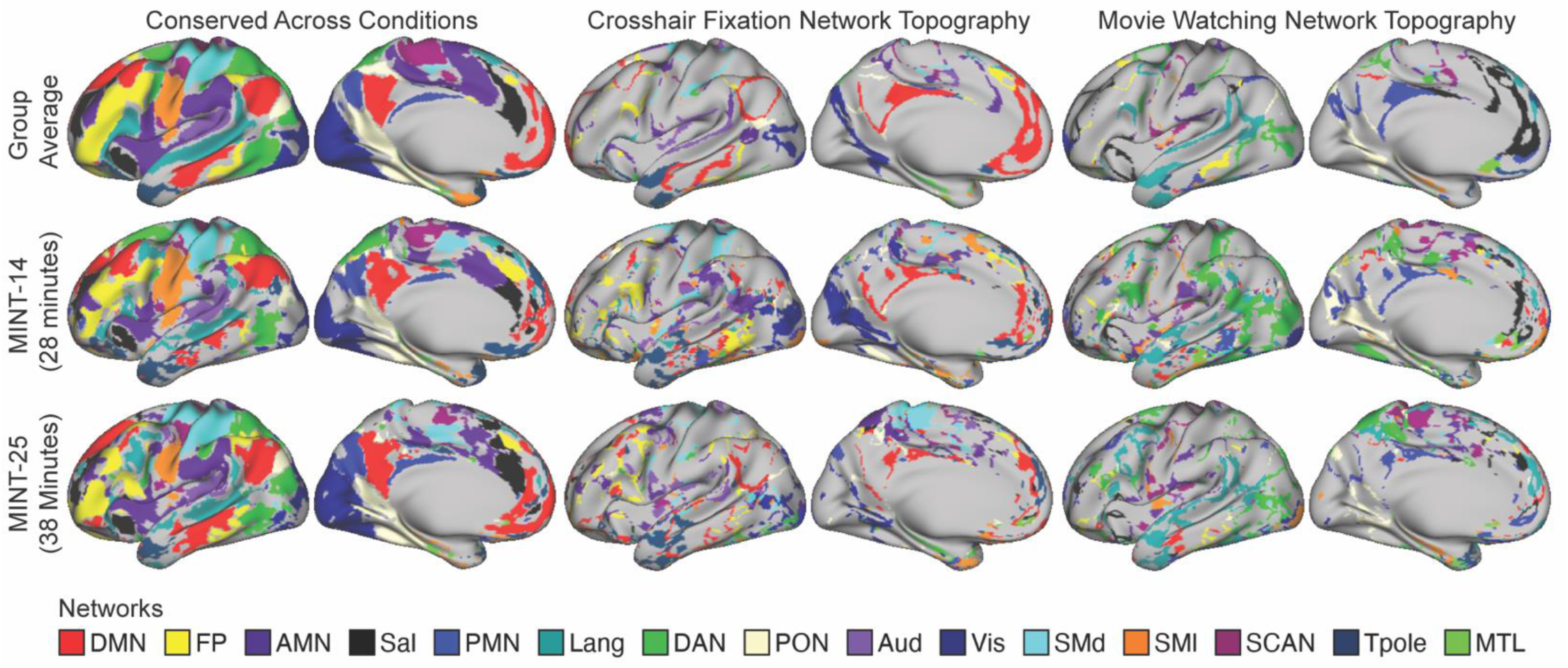
Common and Specific Functional Network Topography Across Crosshair Fixation and Movie Watching. Functional network topography (vertex-wise network assignments) that were conserved and discrepant when derived from crosshair fixation and movie watching data are visualized for the group average and two exemplar participants with the least (28 minutes per-condition; MINT-14) and greatest (38 minutes per-condition; MINT-25) low-motion data.

We next considered these variations in functional network topography and whether they were primarily driven by boundary changes at network borders (‘border shifts’) or could exist at more distal spatial locations between conditions (‘ectopic intrusions’). This analysis follows terminology and parameterization of a previously developed method for characterizing functional network topography variations between individuals to a group average baseline (between-individual approach) (Dworetsky et al., 2024) to instead characterize functional network topography between movie watching to a crosshair fixation baseline (within-individual approach). Using these criteria, the vast majority of movie watching effects on functional network topography in the MINT precision data were border shifts, accounting for 92–98% of the reassigned cortical surface area, while ectopic intrusions accounted for 1–8% of the reassigned cortical surface area across participants. Density maps further show spatial predominance of border shifts across the cortical surface. Border shifts, which were relatively larger in spatial extent, were most predominant in superior temporal gyrus, temporal parietal junction, posterior parietal cortex, and medial prefrontal cortex (Figure 4A). Ectopic intrusions, while relatively smaller in spatial extent, clustered in temporal parietal junction, medial prefrontal cortex, and dorsal anterior cingulate (Figure 4A). These patterns of border and ectopic variants were reproducible across independent data sessions (Figure S6). Collectively, these analyses reveal that the global structure of functional network topography was largely conserved across movie watching and crosshair fixation, evidenced by most reassignments manifesting as border shifts, with minimal reassignments at more distal spatial locations between conditions.

**Figure 4.**
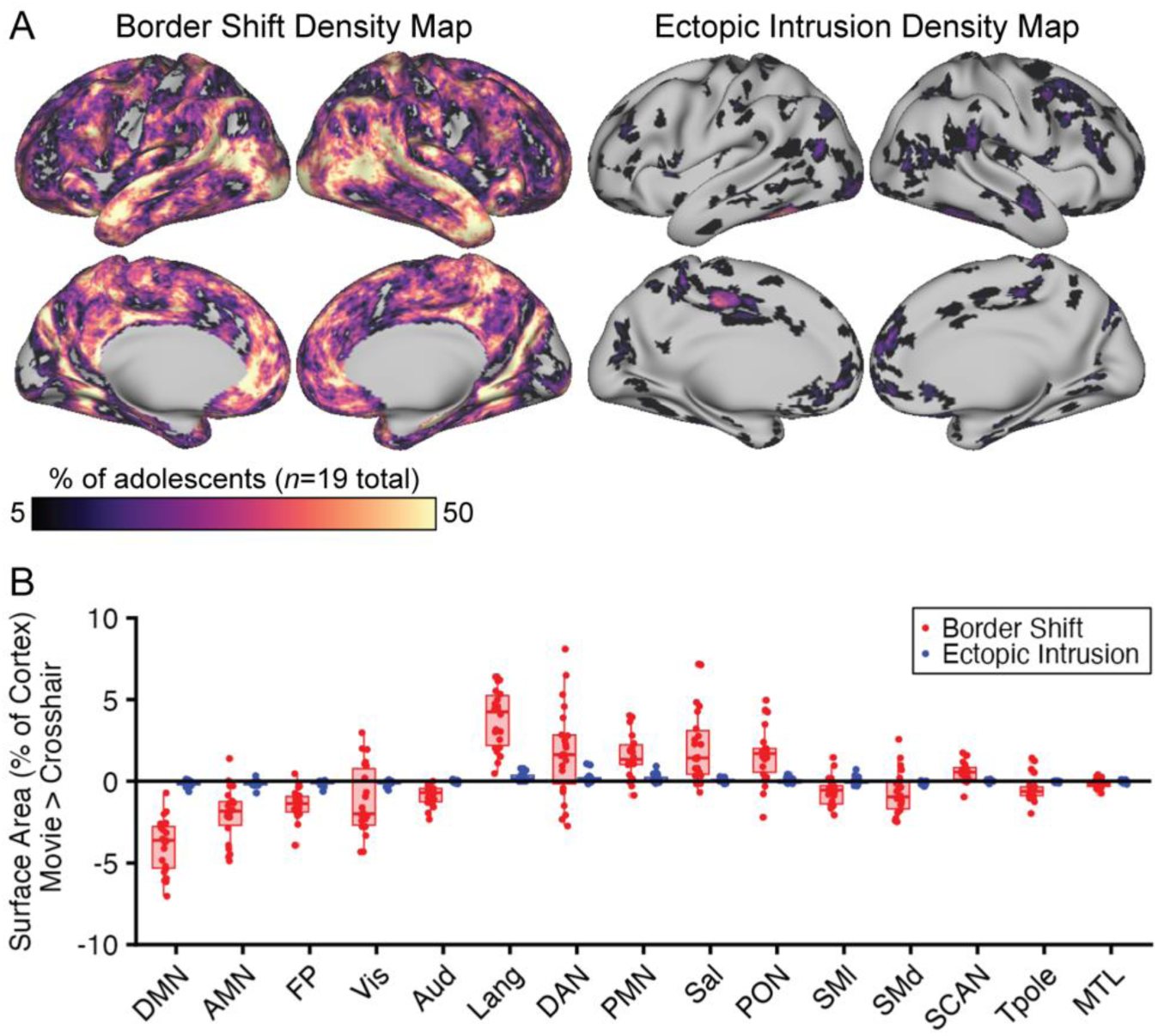
Movie Watching Effects on Functional Network Topography Are Primarily Driven by Boundary Changes at Network Borders. Variations in large scale functional networks per-participant were characterized as border shifts or ectopic intrusions by comparing network assignments during movie watching to participants’ own crosshair fixation baseline following prior terminology and parameterization (Dworetsky et al., 2024). A) Density map showing the proportion of adolescents with border shift or ectopic intrusion at each individual vertex. Border shifts had greater spatial extent across the cortical surface with clusters in superior temporal gyrus, temporal parietal junction, posterior parietal cortex, and medial prefrontal cortex. Ectopic intrusions, relative to border shifts, were smaller in spatial extent and clustered in temporal parietal junction, medial prefrontal cortex, and dorsal anterior cingulate. B) Network reassignments between movie watching and crosshair fixation were primarily border shifts (red) over ectopic intrusions (blue) with the greatest proportion of border shifts and ectopic intrusions observed in networks with the largest variations in intra- and inter-network functional connectivity between viewing conditions (Figure S14).

### Shifting Network Borders During Movie Watching Produce Variable Cortical Network Surface Area Estimates

We next examined whether these border shifts could compound into diverging estimates of cortical network size across viewing conditions. We derived per-participant estimates of cortical network size (% of total cortical surface area in mm^2^, summed across vertices assigned to a network) for crosshair fixation and movie watching data which were compared using linear mixed effects models (see methods for details). Network cortical surface area significantly associated with viewing condition (movie watching > crosshair fixation) for 12/15 canonical networks considered. The largest effects included increased cortical surface area for language network (*β* = 1.61, *pFDR* < 0.001), parietal memory network (*β* = 1.37, *pFDR* < 0.001), and salience network (*β* = 1.24, *pFDR* < 0.001) and decreased cortical surface area for default mode network (*β* = - 1.67, *pFDR* < 0.001) and action mode network (*β* = -1.21, *pFDR* < 0.001) (Figure 5, Table S4). Movie effects in additional networks include increased cortical surface area of dorsal attention network (*β* = 1.04, *pFDR* < 0.01) and parietal occipital network (*β* = 1.12, *pFDR* < 0.001) and decreased cortical surface area of visual network (*β* = -0.81, *pFDR* = 0.010) and auditory network (*β* = -1.06, *pFDR* < 0.001) relative to crosshair fixation (Figure S7, Table S4). These movie watching effects on network cortical surface area estimates were reproducible across independent sessions in the MINT precision data (Figure S8) and there was no interaction between movie watching and functional data quality (mean framewise displacement) for predicting network cortical surface area (Figure S9). See also session to session stability of cortical network size estimates for both movie watching and crosshair fixation (Figure S10).

**Figure 5.**
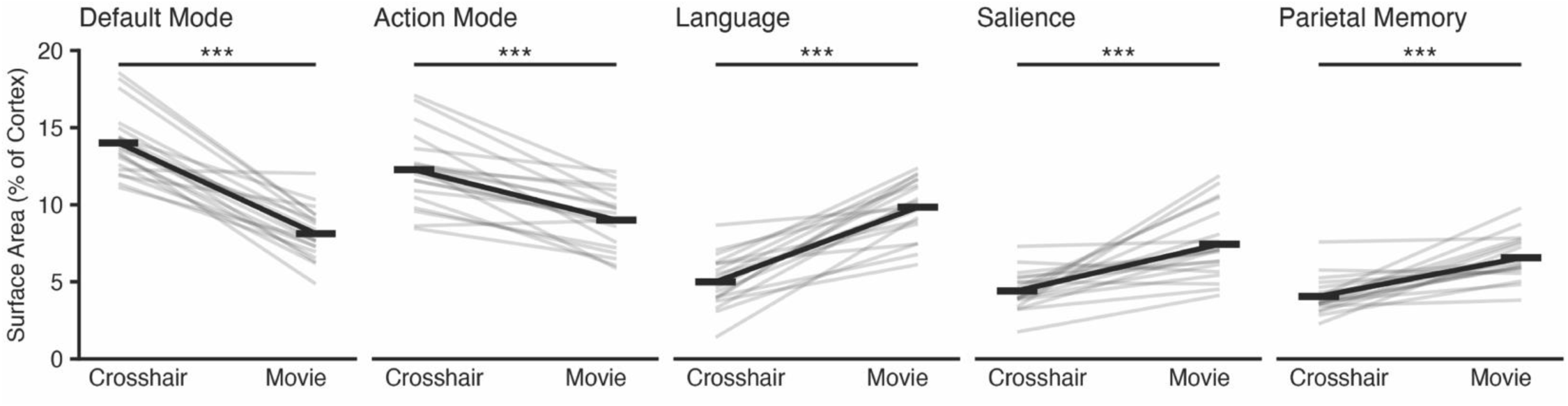
Movie Watching Effects on Cortical Network Surface Area Estimates. Cortical network surface areas by viewing condition are visualized for individual participants (grey lines) alongside the average across participants by condition (black lines). Condition effects on cortical network surface area with high consistency across individual participants (% of total sample) include decreased cortical surface area of default mode (100%) and action mode (95%) networks and increased cortical surface area of language (100%), parietal memory (95%), and salience (89%) networks. Statistical significance was determined using separate linear mixed effects models for each network, with cortical surface area as the outcome and a main effect of viewing condition (movie watching > crosshair fixation) while covarying for fixed effects of age, sex, and head motion (mean framewise displacement, mm) and random subject intercepts. False discovery rate correction was applied across the 15 network linear mixed effects models. ***: *p_FDR_* < 0.001. Descriptive and inferential statistics for cortical network surface area by viewing condition are available in the supplemental materials (Table S4).

### Relative Magnitude of Group, Individual, and Viewing Condition Effects on Network Topography

We next looked to quantify the movie watching effect relative to condition-invariant group and individual effects on functional network topography to further contextualize movie watching as an alternative acquisition state for assessing individual differences. We therefore contextualized condition, individual, and group effects by calculating the increased similarity of network assignments vectors (Dice-Sørensen Coefficients) paired by these features, which were normalized relative to the maximal similarity, to get a normalized relative magnitude of each additive ‘effect’ as applied previously to functional connectivity (Gratton et al., 2018) (see methods for further details). The largest effect on whole-cortex functional network assignments was baseline similarity across adolescents and viewing conditions (the shared ‘group’ effect) accounting for 61% of the normalized relative variance (Figure 6A). There was a minimal increase over this group similarity for functional brain networks from the same viewing condition but different individuals (the ‘condition’ effect, 5% relative normalized variance) representing a shared condition response across individuals (Figure 6A,B). There was a large jump in similarity for functional brain networks from the same individual across viewing conditions, representing trait-like individual differences that persist across conditions (the ‘individual’ effect, 19% of the relative normalized variance) (Figure 6A,B). Functional brain networks from the same individual and same viewing condition were the most similar, representing individual variability in the condition response (the ‘individual by condition effect’, 14% of relative normalized variance) (Figure 6A,B). These observations demonstrate that functional network topography in clinical adolescents are predominantly driven by common underlying properties of the group (61%) and trait-like individual differences (19%). See also relative effect magnitudes for functional connectivity as in previous work (Gratton et al., 2018) (Figure S11).

**Figure 6.**
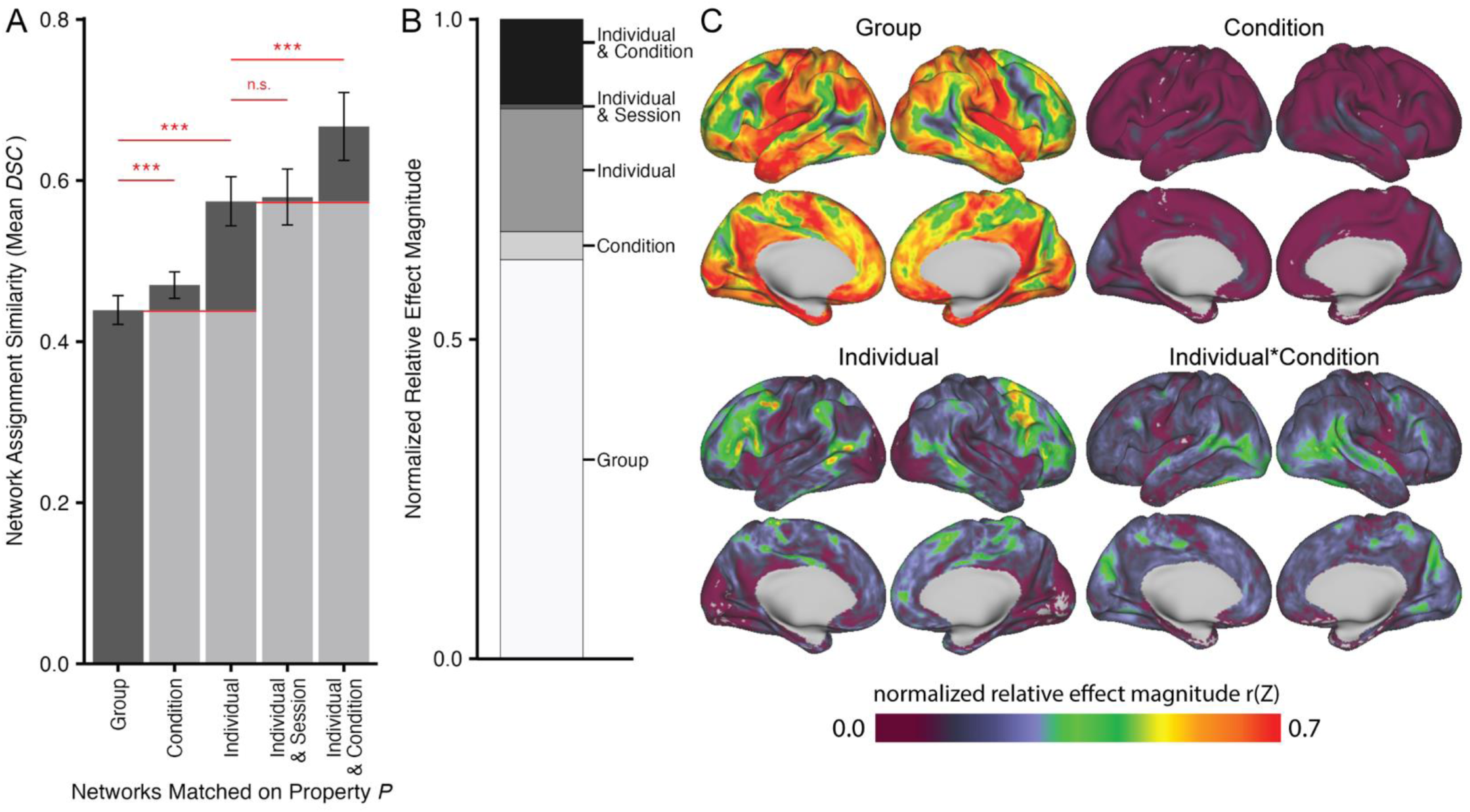
Magnitude and Spatial Distributions of Group, Individual, and Viewing Condition Effects on Functional Network Assignments. A) Mean similarity of whole-cortex network assignment vectors at the level of group (different adolescents, different conditions), condition (different adolescents, same condition), individual (same adolescent, different conditions), individual by session (same adolescent, different conditions, but the same session), and individual by condition (same adolescent, same condition). B) The improved similarity of whole-cortex network assignment vectors was quantified relative to the maximum similarity (here, the individual by condition similarity) to get normalized relative magnitude of each additive ‘effect’ following the approach of (Gratton et al., 2018). The largest contributors to whole cortex network assignments were group similarity and individual differences. C) The same similarity analysis at the whole cortex level was repeated at each individual vertex (each row of the vertex-to-network similarity matrix) to quantify the normalized relative magnitude of each effects across the cortical surface. See also spatial distributions of group, condition, and individual by session effects on a narrower color range (Figure S12).

### Cortical Spatial Distributions of Group, Individual, and Viewing Condition Effects on Network Assignments

To examine whether group, individual, and viewing condition effects had common or distinct spatial distributions across the cortical surface, we applied a similar approach at individual vertices by calculating similarities between template matching intermediate vertex-to-network similarity vectors (15 network similarities for each vertex *i*) (see methods). Group effects (baseline similarity across different adolescents and different viewing conditions) were high in several regions including somatosensory cortex, primary auditory cortex, and insula (Figure 6C). Individual effects (increased similarity over the group effect for the same adolescent across viewing conditions) in contrast were highest in dorsolateral prefrontal cortex, posterior parietal cortex, and temporal parietal junction (Figure 6C), suggesting these regions demonstrate trait-like individual variation detectable across conditions. Condition effects (similarity between different adolescents but the same condition) were relatively minimal in contrast to group and individual effects, but were highest in primary and secondary visual cortex and secondary auditory cortex (Figure 6C) which likely represents common low-level stimulus processing. Individual by condition effects (increased similarity over the individual effect for observations from the same individual and same condition) were highest in regions that support secondary stimulus processing including temporoparietal junction, frontal eye fields, secondary visual cortex, and posterior cingulate cortex (Figure 6C). Individual by session effects (similarity within adolescents, for different conditions but the same session) were generally low across the brain (Figure S12). Cortical spatial distributions of group, condition, and individual by session effects on custom color scales are visualized in the supplement (Figure S12).

Visual inspection of group, individual, and individual by condition effects across the cortical surface (Figure 6C) suggest these effects have variable spatial distributions over three clinically relevant regions of interest. The group effect appears highest in somatomotor cortex (SM), including supplementary motor area, which is a neuromodulation site for obsessive compulsive disorder (Hawken et al., 2016; C. Conelea et al., 2024) and tic-related disorders (Wu et al., 2014; C. Conelea et al., 2023), the individual effect appears highest in dorsolateral prefrontal cortex (DLPFC) which is a neuromodulation site for mood disorders including major depressive disorder (Fox et al., 2013; Cash et al., 2021; Cole et al., 2022), and the individual by condition effect appears to be highest in temporal parietal junction (TPJ) which is a clinically relevant neuromodulation region for schizophrenia and autism (Ahmad et al., 2021; Dabiri et al., 2022).

To confirm these observations we examined whether the spatial effect distribution maps (Figure 6C), derived using template matching intermediate vertex-to-network similarity vectors, accurately predict varying contributions of group, individual, and condition effects on functional network topography across clinically relevant regions. We derived surface masks for SM, DLPFC, and TPJ using independent data from Neurosynth (Yarkoni et al., 2011) (Figure 7A) then repeated the same similarity analyses on winner-take-all network assignment vectors within each region of interest. In SM there was a larger relative contribution of group (70% of maximum similarity) over condition (2%), individual (19%), or individual by condition (9%) effects (Figure 7B). In DLPFC, though the largest relative contributor was group (50% of maximum similarity) there was a larger individual effect (34%) compared to other regions of interest (Figure 7C). In TPJ there was a larger relative contribution of the condition effect compared to other regions (7% of maximum similarity) and the individual (22%) and individual by condition (23%) effects were similar in magnitude (Figure 7D).

**Figure 7.**
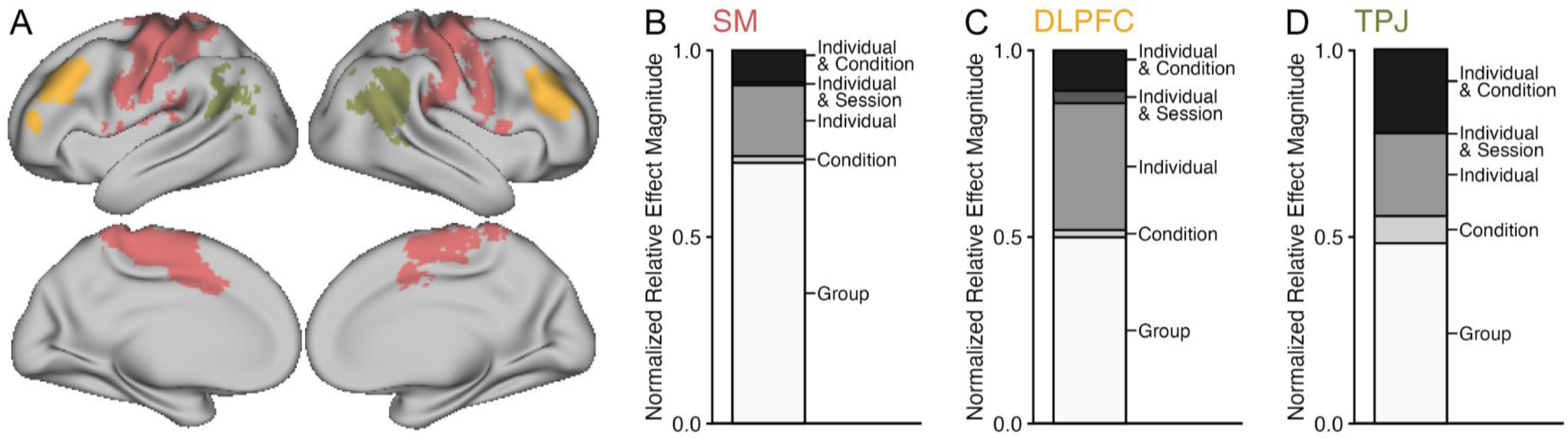
Group, Individual, and Viewing Condition Effects on Functional Brain Network Topography in Clinically Relevant Regions of Interest. A) Independent data from the neurosynth database (Yarkoni et al., 2011) were used to derive masks for three clinically relevant regions of interest which were somatomotor cortex (SM), dorsolateral prefrontal cortex (DLPFC), and temporal parietal junction (TPJ). Region of interest masks spanned the left and right cortical hemispheres but are visualized on the right hemisphere. B) SM network assignments were predominantly driven by stable group effects, and these group effects were higher in SM compared to other regions of interest (Figure 6C,D) or the whole cortex (Figure 6B). C) DLPFC network assignments had a larger relative contribution of individual effect, individual similarity that persists across viewing conditions, compared to other regions of interest (Figure 6C,D) or the whole cortex (Figure 6B). D) TPJ network assignments had a larger relative contribution of condition and individual by condition effects compared to other regions of interest (Figure 6C,D) or the whole cortex (Figure 6B). *SM:* somatomotor cortex, *DLPFC:* dorsolateral prefrontal cortex, *TPJ:* temporal parietal junction.

### Replication of Key Results in Healthy Brain Network Consortia Data

In our final set of analyses we examined the replicability of key results using independent consortia data from the Healthy Brain Network (n = 466, age = 6–22 years). In the HBN consortia data we replicate observations that most viewing condition effects on network assignments manifested as boundary changes at network borders (border shifts, 84-100% of the reassigned cortical surface area in mm^2^), while spatially distinct variations >3.5mm were uncommon (ectopic intrusions, 0-16% of the reassigned cortical surface area in mm^2^). In the HBN data we further replicate that border shifts were greater in spatial extent while being most prominent in superior temporal gyrus, temporal parietal junction, medial prefrontal cortex, and dorsal anterior cingulate (Figure 8A). Ectopic intrusions in the HBN consortia data were relatively smaller in spatial extent compared to ectopic intrusions in the MINT precision data; however, the ectopic intrusions across datasets showed small clusters in temporal parietal junction, medial prefrontal cortex, and dorsal anterior cingulate (Figure 8A). The HBN consortia data further showed replicable network-specific patterns of border and ectopic variants, with the greatest proportion of both border shifts and ectopic intrusions involving spatial contraction of default mode and action mode networks and spatial expansion of language and dorsal attention networks (Figure 8A).

**Figure 8.**
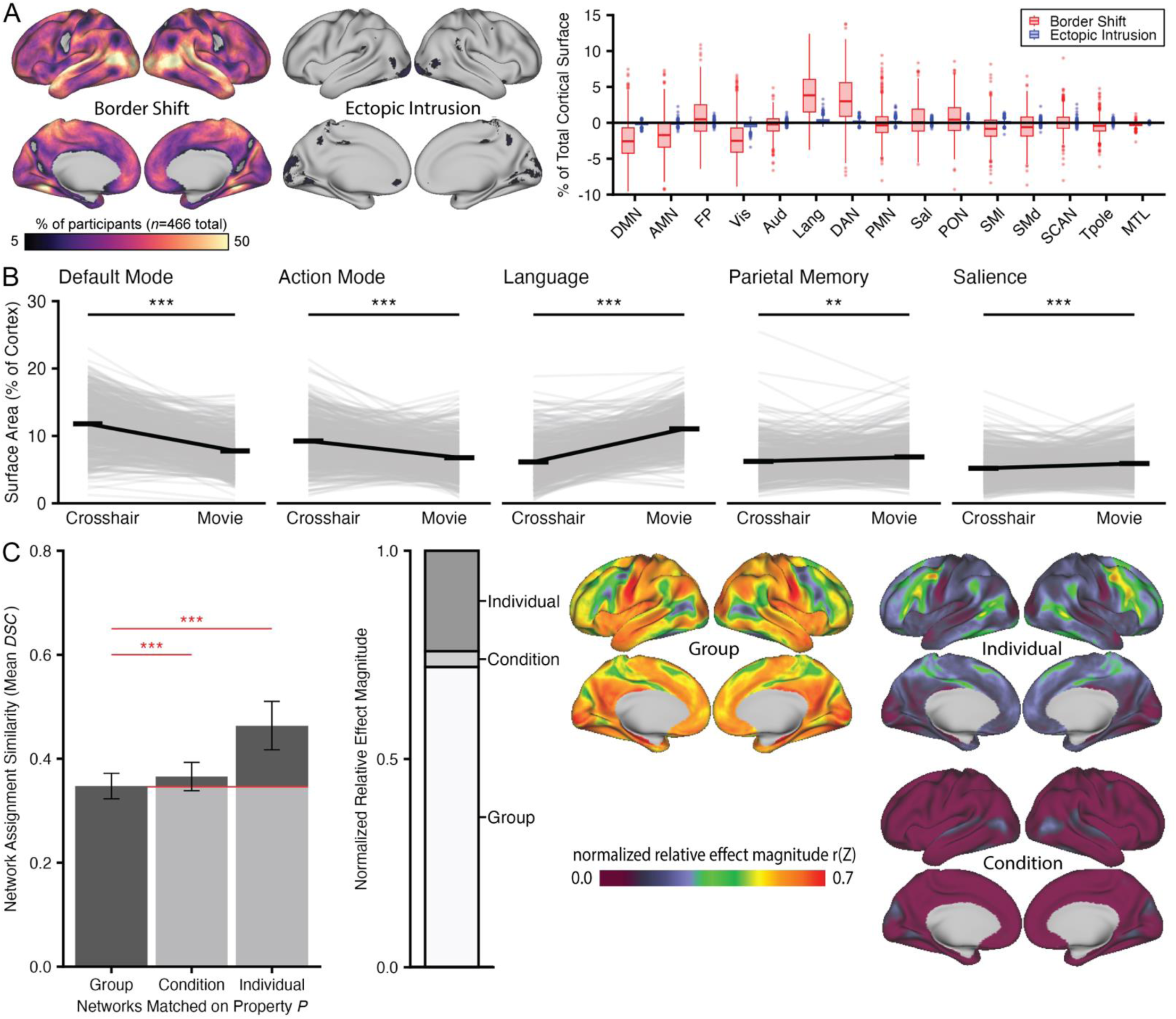
Replication of Key Results in Independent Healthy Brain Network Consortia Data. A) In the HBN consortia data (*n*=466), replicating patterns observed in the MINT precision data, border shifts were relatively larger in spatial extent clustering in superior temporal gyrus and temporal parietal junction. Ectopic intrusions were smaller in spatial extent in the HBN consortia data, while having small clusters in temporal parietal junction, dorsal anterior cingulate, and medial prefrontal cortex. B) Replicable effects of condition on cortical network surface include decreased cortical surface area of default mode and action mode networks and increased cortical surface area of language, salience, and parietal memory networks alongside a more distributed pattern of replicable effects (Figure S13). C) In the HBN consortia data, as in the MINT precision data, functional network assignments were predominantly driven by group effects (clustering in somatomotor cortex) and individual effects (clustering in dorsolateral prefrontal cortex) above a shared condition-related response across participants (the ‘condition’ effect, clustering in superior temporal gyrus). ***: *p_FDR_* < 0.001.

In the HBN consortia data we additionally replicated results demonstrating directionality of movie watching effects on cortical network surface area (% of total cortical surface area in mm^2^) which were decreased cortical surface area estimates of default mode (*β* = -0.95, *pFDR* ≤ 0.001) and action mode networks (*β* =-0.69, *pFDR* ≤ 0.001) and increased cortical surface area estimates of language (*β* = 1.27, *pFDR* ≤ 0.001), parietal memory (*β* = 0.98, *pFDR* ≤ 0.001), and salience networks (*β* = 0.19, *pFDR* = 0.002) (Figure 8B). Additional replicable effects were also seen for decreased cortical surface area estimates of visual network (*β* =-1.15, *pFDR* ≤ 0.001) and auditory network (*β* =-0.41, *pFDR* = 0.025) and increased cortical surface area estimates of parietal occipital network (*β* = 0.28, *pFDR* ≤ 0.001) and dorsal attention network (*β* =0.39, *pFDR* ≤ 0.001) (Figure S13). Descriptive and inferential statistics for viewing condition effects on cortical network surface area in the HBN consortia data are fully summarized in the supplemental materials (Table S5).

Finally, in the HBN consortia data we replicate observations that large-scale functional network assignments are predominantly driven by shared common underlying features of the group (the ‘group’ effect) and trait-like individual variability (the ‘individual’ effect) with a relatively minimal contribution of viewing condition (the ‘condition’ effect) (Figure 8C). When examining spatial distributions of these effects across the cortical surface, as in the MINT precision data, group effects clustered in somatomotor cortex, individual effects clustered in dorsolateral prefrontal cortex and temporal parietal junction, and condition effects (though relatively smaller in magnitude and spatial extent than group and individual effects) clustered in superior temporal gyrus and temporal parietal junction (Figure 8C). We note that the HBN data is collected at a single session and therefore lacks the test-retest design necessary to derive the individual by condition effect (i.e., increased same individual similarity for functional network topography derived from the same viewing condition across data sessions) so the individual by condition effect could not be tested for replicability.

## DISCUSSION

We assessed individual large-scale functional network topography derived from movie watching and crosshair fixation using precision functional data collected in a cohort of adolescents with transdiagnostic psychiatric conditions with key results reproduced across independent data sessions and replicated using independent consortia data from the Healthy Brain Network. We demonstrate that the global structure of large-scale functional networks is generally conserved between conditions with most movie effects manifesting as border shifts between neighboring networks. Though the general global structure of functional network topography was conserved, choice of viewing condition produced notable differences in cortical surface area estimates which were most prominent for movie-associated functional networks. Despite these observed differences, functional data collected during both movie watching and crosshair fixation can be used to derive individual functional brain network topography with similar individual-specificity and good test-rest reliability. Movie effects on individual functional brain network estimates were notably minimal for two frequent clinical neuromodulation targets (dorsolateral prefrontal cortex and somatomotor cortex) while being more prominent in a third (temporal parietal junction) highlighting region of interest considerations for movie watching as an alternative acquisition to crosshair fixation. We will synthesize these results to suggest where movie watching could be considered as an alternative to crosshair fixation for individual functional network topography estimation, particularly for pediatric and clinical samples that may struggle with conventional resting-state data collection using crosshair fixation (Emerson et al., 2015; Long et al., 2017; Espenhahn et al., 2021).

### Global structure of large-scale functional network topography is conserved across conditions

The global structure of large-scale functional network topography was generally conserved when estimated from movie watching compared to crosshair fixation. Indeed, most of the variable cortical surface area (92-98%) between conditions were border variants that reflect shifting boundaries between neighboring networks while the global structure of network ‘core’ spatial locations was largely conserved. These observations are consistent with the perspective that most viewing condition effects are driven by relatively subtle underlying functional connectivity variations (see Figure S14). A smaller subset of network reassignments during movie watching were ‘ectopic’ intrusions involving reassignments between more distal network territories (>3.5mm) following parameterization in prior work (Dworetsky et al., 2024). Functional networks most likely to form ectopic intrusions (language, salience, and parietal memory networks) had underwent the greatest increase in intra- or inter-network functional connectivity (Figure S12). Therefore, though most movie effects on large-scale functional networks can likely be attributed to subtle variations between neighboring networks within a conserved global structure, a small subset of ‘ectopic intrusions’ were detected for movie-positive functional networks.

### Movie watching and crosshair fixation are more distinct when considering cortical network size

While these results suggest generally conserved global structure of large-scale functional networks across viewing conditions, we further report notable viewing condition effects when deriving estimates of cortical network size. The largest effects of movie watching on cortical network size were decreased surface area estimates of default mode and action mode networks and increased cortical surface area estimates for language, parietal memory, and salience networks all of which displayed significant changes in connectivity profiles between movie watching and crosshair fixation (Figure S12). Integrating across prior reports and our results, we suggest that movie watching can be expected to decrease cortical surface area estimates for movie-negative networks while increasing cortical surface area estimates for movie-positive networks. We also note that while differences between movie watching and crosshair fixation conditions have minimal effects on network ‘cores’, it appears that border shifts between the two conditions compound into substantive differences in the apparent relative size of these networks when assigning network identities with a winner-take-all heuristic. Identifying network labels with a winner-take-all heuristic, where each vertex is given a single network identity, may therefore amplify subtle variations in functional connectivity into larger observed effects when considering cortical network size.

### Movie watching for individual functional network estimation in clinically-relevant neuromodulation targets

Movie watching may be particularly beneficial for precision functional mapping in pediatric clinical cohorts with an indication for individual brain network guided neuromodulation. Assessing movie watching as a suitable alternative acquisition state requires consideration of the relative magnitude of viewing condition effects across clinically relevant regions of interest and determining whether movie watching preserves individual differences in network topography. We therefore examined the relative effects of group (stable between person differences), individual (individual differences that persist across conditions), and condition (movie watching vs. crosshair fixation) on functional network topography in three clinically relevant regions of interest.

A large individual effect, individual differences in network topography that persist across viewing conditions, was observed in dorsolateral prefrontal cortex (DLPFC) which is a treatment target for mood disorders including major depressive disorder (Fox et al., 2013; Cash et al., 2021; Cole et al., 2022). Prior work in adults suggests DLPFC is one of the most individualized brain regions (Mueller et al., 2013; Fox et al., 2013; Ladwig et al., 2026) and that neuromodulation applied to DLPFC benefits from defining individual-specific target sites (Fox et al., 2013; Cash et al., 2021; Cole et al., 2022). These results demonstrate that the same individual principles observed in adults are translated to adolescents, which suggests that accounting for individual functional network variability will continue to be particularly critical for DLPFC. The relative stability of individual network features in DLPFC across viewing conditions indicates that movies can be considered reasonably comparable for estimating individual network boundaries in this region. The context-independent individualization of DLPFC is particularly relevant considering movies have been shown to evoke functional alterations in prefrontal regions (Vanderwal et al., 2015; Kringelbach et al., 2023), supporting arguments of trait-like functional network topography in individuals (Seitzman et al., 2019; Kraus et al., 2021; Dworetsky et al., 2024; Ladwig et al., 2026).

A large group effect, high stability of network topography across individuals regardless of viewing condition, was observed in somatomotor cortex. Somatomotor areas, including supplementary motor area, are neuromodulation targets for psychiatric conditions including obsessive compulsive disorder (Hawken et al., 2016; C. Conelea et al., 2024) and tic-related disorders (Wu et al., 2014; C. Conelea et al., 2023). A large group effect in somatomotor cortex suggests this region is relatively less individual-specific and, when necessary, the use of group defined network boundaries will be more likely to represent individual network boundaries for somatomotor cortex compared to other brain regions. However, the group effect did not completely account for all variance in somatomotor cortex, which suggests caution in assuming strict group to individual generalizability of functional organization and that individual identification likely remains useful.

A large individual by condition effect, individual differences that vary across viewing conditions, was observed in TPJ which is a clinically relevant region for conditions including autism and schizophrenia (Ahmad et al., 2021; Dabiri et al., 2022). These results suggest that for TPJ, it’s both essential to determine network boundaries at the individual level and be cognizant of potential confounding viewing condition effects. Given individual variability in the viewing condition response, future investigations considering TPJ may benefit from collecting fMRI across both crosshair fixation and movie watching acquisitions to fully characterize individual differences in this region.

### Recommendations for estimating individual large-scale functional network topography with movies vs. crosshair

We examined movie watching and crosshair fixation for individual functional network topography estimation across multiple spatial scales, derivative measures, and datasets. Here, we synthesize these findings to provide several key insights for future study planning. We identified that movies reduced head motion with the greatest benefits observed for younger children which replicates prior reports (Greene et al., 2018; Frew et al., 2022). Integrating our results with prior work, we therefore suggest that future studies recruiting pediatric participants could consider movie watching to improve data quality. Improved data quality does not necessarily mean that movie watching and crosshair fixation are interchangeable for all scientific questions or clinical goals. The choice of viewing condition should also consider if the specific measures of interest differ across viewing conditions as discussed next.

#### Network ‘cores’ vs cortical network size

Movie watching and crosshair fixation are likely more comparable for analyses deriving network ‘core’ spatial locations, which may be of clinical interest for individual precision neuromodulation which aims to identify network centroids for optimal coil placement (Lynch et al., 2022). While this work suggests that the global structure of functional network core regions is generally conserved in a cohort of adolescents most likely to benefit from individual brain network guided neuromodulation, we caution that the efficacy of neuromodulation protocols across movie watching and crosshair fixation remains unknown until directly tested.

In contrast movie watching and crosshair fixation are less comparable for studies concerned with cortical network size and multi-network border ‘encroachment’ patterns (Lynch et al., 2024; Vaghi et al., 2025). The networks with the most variable cortical surface area estimates between movie watching and crosshair fixation, particularly salience and default mode, are relevant to investigations considering associations between network size and psychopathology. Specifically, prior work has associated salience network expansion and encroachment into default mode or frontoparietal networks with symptom burden in adults with major depressive disorder (Lynch et al., 2024) and obsessive compulsive disorder (Vaghi et al., 2025). Investigators interested in characterizing associations between network size and behavior or psychopathology will therefore have to be more judicious when selecting between movie watching and crosshair fixation; and whether functional network topography derived from movie watching or crosshair fixation is superior for predicting individual differences has not been tested. We further suggest caution when comparing across published reports or harmonizing datasets using different viewing conditions (movie watching vs. crosshair fixation) with the intention of examining cortical network size unless confounding viewing condition effects are properly considered.

#### Network topography vs. dynamics and connectivity

We examined movie watching and crosshair fixation for deriving individual estimates of large-scale functional network topography (winner-take-all vertex network assignments). This study was designed to provide practical guidance for clinical applications by collecting feasible levels of data in a cohort of adolescents most likely to benefit from movie watching for individual brain network guided neuromodulation. We demonstrate that, depending on the level of analysis, movies can be considered as an alternative to crosshair fixation for individual functional network identification with static functional connectivity. Owing to the focus on individual network estimation this work primarily focused on functional network topography, rather than in-depth analysis of functional connectivity changes. While our analysis of functional network connectivity is aligned with prior work (c.f., (Emerson et al., 2015; Vanderwal et al., 2017)) we acknowledge additional reports of movie effects on measures beyond functional network topography including intersubject correlation (Hasson et al., 2004, 2008; Mochalski et al., 2024) and brain state transitions (van der Meer et al., 2020; Song et al., 2023; Hughes et al., 2025). Investigators should therefore consider the breadth of further analyses beyond functional network topography and potential effects of movie watching when selecting between viewing conditions.

Similarly, as our focus was on practical guidance for functional network topography estimation, we focused on movie watching as an acquisition “state” rather than analyzing the dynamic stimulus features and cognitive, affective, or social processes that may give rise to these effects. Movies are highly complex, irregular, and dynamic across high-level narrative and low-level stimulus variations (Vanderwal et al., 2019) which evoke connectivity alterations when compared to crosshair fixation across sensory, cognitive control, attention, and association networks (Bottenhorn et al., 2018). It would be of interest for future work to consider functional signal properties and potential mechanisms that underlie the effects of movie watching on functional network topography reported here. This may include, for example, whether functional network topography effects are driven by global or local signal variations (e.g., luminance changes influencing visual network and narrative changes influencing default mode network) or whether functional network topography estimates vary across the course of movie viewing. Answering these questions would require a more stringent event-related design which was not considered here. In contrast, analyses focused on functional network topography estimation and primarily interested in selecting an acquisition state may wish to consider movie watching as an alternative to crosshair fixation noting individual, regional, and analytic variability.

## Conclusions

Together, these findings demonstrate that precision functional mapping (PFM) in adolescents across transdiagnostic categories yields similarly individual-specific and test-retest reliable functional brain network topography estimates when derived from crosshair fixation and movie watching acquisitions. Analyses across multiple functional network indicators and levels of spatial scale reveal notable and reproducible differences, but a high degree of consistency across conditions. This work demonstrates instances where movie watching could be considered as an alternative to crosshair fixation for deriving individual large scale functional network topography while establishing key considerations for individual variability, regional differences, and condition-dependent effects in certain types of analyses. The results here can help guide broader implementation of movie watching for individual brain network mapping in pediatric and clinical populations where traditional crosshair fixation may not be feasible.

## METHODS

### Mapping Individual Networks Transdiagnostically (MINT) Precision Dataset

#### Participants Overview

We aimed to enroll a sample of youth with neurodevelopmental and psychiatric disorders for which neuromodulation interventions are currently available or are being tested in ongoing research that could be enhanced with precision imaging methods. In the United States, ∼20% of children experience a psychiatric disorder in a given year (Sappenfield et al., 2024). The most prevalent disorders diagnosed in youth are attention-deficit hyperactivity disorder, anxiety, depression, autism spectrum disorder, and Tourette Syndrome (Bitsko et al., 2022). Currently available, evidence-based interventions for these disorders include psychotherapy, pharmacotherapy, and, in some cases, non-invasive brain stimulation, yet efficacy is limited, with many meta-analyses showing moderate effects (e.g., (Weisz et al., 2006; Correll et al., 2021)). Individual differences in brain structure and function are likely to play a role in disease trajectories and treatment response (Shin et al., 2013), particularly for emerging treatments focused on targeted neuromodulation (C. A. Conelea & Lieske, 2026). To increase the likelihood that this methods development study could be readily applicable for clinically orientated research, we elected to focus on the target population of need.

Participants were recruited from research registries, digital advertising, flyering, and distribution of study information to local healthcare providers. Inclusion criteria were as follows: 1) Age 7-17 years (inclusive) at the time of enrollment; 2) Meet DSM-5 criteria for a current diagnosis of depressive disorder (including major depressive disorder and disruptive mood dysregulation disorder), anxiety disorder (including generalized Anxiety Disorder, Social Anxiety Disorder, Specific Phobia, Separation Anxiety Disorder), obsessive compulsive disorder, tic disorder/Tourette Syndrome, autism spectrum disorder, attention deficit hyperactivity disorder (ADHD) or adjustment disorder, as established via records and/or confirmation with the The Mini International Neuropsychiatric Interview-7 (Sheehan et al., 1998); 3) Participants taking psychotropic medications were included if the medications were stable for 6 weeks with no anticipated changes for the duration of their study participation (ie., through the second MRI visit); 4) English fluency; 5) Right-handedness. Exclusion criteria were 1) Inability to undergo MRI; 2) Previous or current diagnosis of psychosis, cognitive disability, or bipolar disorder, 3) Active substance use or dependence disorder; 4) Active suicidality; 5) Left handedness; or 6) any other medical or psychiatric condition deemed serious or contraindicated for any study procedures by the study team. This study was approved by the Institutional Review Board at the University of Minnesota. Consent was obtained from a parent or legal guardian and assent was obtained from children prior to data collection. If a minor turned 18 during the course of their participation, they were re-consented as an adult.

#### Behavior and Assessments

A preliminary eligibility assessment was completed over the phone with interested parents. Those deemed potentially eligible completed a clinical assessment visit to confirm eligibility and complete the assessment battery, which included: 1) a demographics form to enable sample characterization; 2) the Mini International Neuropsychiatric Interview-7 (Sheehan et al., 1998), a structured diagnostic interview for the most common childhood DSM-5 and ICD-10 psychiatric disorders; 3) the Spence Children’s Anxiety Scale-Parent and Child Report (SCAS) (Spence, 1998; Spence et al., 2003), a measure of anxiety symptom severity; 4) the Children’s Depression Inventory (CDI) (Kovacs, 1992).

#### Magnetic Resonance Imaging

Magnetic resonance imaging (MRI) data were collected from participants at each of two independent sessions less than one month apart (days between scanning sessions; mean = 6, standard deviation = 5, range = 1–18) at the Masonic Institute for the Developing Brain. MRI data were collected using a 3T Siemens Prisma with a 32 channel headcoil. Each MRI session included a T1 (TR = 2.50s, TE = 2.88ms, flip angle = 8°; voxels = 1mm^3^) and T2 (TR = 3.20s, TE = 5.65s, flip angle = 120°, voxels = 1mm^3^) anatomical scan; AP and PA spin echo field maps (single echo, single-band, TR=13.70s, TE=2.90ms, flip angle = 90°); and four functional runs of T2* multi-echo, multi-band, echo-planar imaging (TR = 1.76s; TEs = 14.2, 38.94, 63.66, 88.40ms; MB factor = 6; IPAT = 2; flip angle = 68°; voxels = 2mm^3^). Functional MRI (fMRI) at each session were collected for movie watching (concatenated *Pixar* movie shorts for children) and crosshair fixation (white crosshair, black background). Functional stimuli were presented in the same order across participants. Session 1: movie watching (1x10 minutes; *The Blue Umbrella, Jack-Jack Attack*), crosshair fixation (1x10 minutes), movie watching (1x10 minutes; *The Adventures of Andre and Wally B., Burn-E, Marine Life Interviews*), crosshair fixation (1x10 minutes); Session 2: move watching (1x10 minutes; *Lifted*, *Hawaiian Vacation*), crosshair fixation (1x10 minutes), movie watching (1x10 minutes; *For the Birds, Dug’s Special Mission, Miss Fritter’s Racing Skoool*), crosshair fixation (1x10 minutes).

### Healthy Brain Network (HBN) Consortia Dataset

#### Participants Overview

A full description of the Healthy Brain Network (HBN) consortia data is available elsewhere (Alexander et al., 2017), here we will provide a brief overview of the data used in the present study. The objective of the HBN consortia is to recruit a large-scale sample of children and adolescents in the New York City area that are representative of transdiagnostic neurodevelopmental and psychiatric disorders which present in developmental samples. Participants were therefore recruited through a community-referred recruitment model where advertisements encouraged participation from families concerned about psychiatric symptoms in their children (Alexander et al., 2017).

Participants were recruited using advertisements distributed to community members, educators, local care providers, email lists, and events. Inclusion criteria were as follows: 1) male or female children aged 5-21 years; 2) capacity of parents for informed consent; 3) child English fluency; 4) parent English fluency or parent Spanish fluency alongside availability of Spanish speaking personnel. Exclusion criteria were: 1) cognitive or behavioral impairment (e.g., IQ < 66); 2) acute encephalopathy; 3) a known neurodegenerative disorder (e.g., Huntington’s Disease); 4) hearing or visual impairment without assistive devices; 5) recent diagnosis (past 6 months) of schizophrenia, schizoaffective disorder, or bipolar disorder without treatment; 6) acute manic or psychotic episode without current and ongoing treatment; 7) new onset (past 3 months) suicidality or homicidality without current and ongoing treatment; 8) history of lifetime substance dependence requiring clinical replacement therapy; 10) acute intoxication at any time of study visit. These and further details regarding HBN consortia participant recruitment and inclusion/exclusion criteria are available elsewhere (Alexander et al., 2017).

#### Behavior and Assessments

A preliminary eligibility screening was performed over the phone with interested parents (Alexander et al., 2017). Eligible participants completed the following assessment battery: 1) a demographic survey to enable sample characterization; 2) the Schedule for Affective Disorders and Schizophrenia - Children’s version (KSADS) (Kaufman et al., 1997), a semi-structured interview for deriving clinical diagnoses; 3) the Child Behavior Checklist (CBCL) (Achenbach, 1991), a parent report questionnaire to assess emotional and behavioral problems in children. Where indicated by the KSADS, follow up assessments were administered to confirm suspected diagnoses (for details, see (Alexander et al., 2017)).

#### Magnetic Resonance Imaging

HBN Release 11 data were accessed for the present study (Alexander et al., 2017). We identified participants from Rutgers University Brain Imaging Centre and CitiGroup Cornell Brain Imaging Centre with complete data (n=1377) defined as: 1) a T1 anatomical, 2) complete functional data for crosshair fixation (2 x 5 minute runs; white crosshair on a black background) and movie watching (*The Present*: 1 x 3 minute run and *Despicable Me*: 1 x 10 minute run), 3) accompanying field maps. MRI data were collected on either a 3T Siemen’s Tim Trio (Rutgers University Brain Imaging Center) or Siemens 3T Prisma (CitiGroup Cornell Brain Imaging Center).

Each MRI scan included in the present study consisted of one or more T1 anatomicals collected using protocols from the Human Connectome Project (Van Essen et al., 2012) or Adolescent Brain Cognitive Development Study (Casey et al., 2018); AP and PA spin echo field maps; and four functional runs of T2* echo-planar imaging (TR = 800ms; TE = 30ms; MB factor = 6; flip angle = 31°; 2.4mm^3^ resolution) (Alexander et al., 2017). Functional MRI (fMRI) at each session were collected for crosshair fixation (white cross, black background) and movie watching (*Despicable Me* and *The Present*). Functional stimuli were presented in the same order across participants: crosshair fixation (1x5 minute run), movie *Despicable Me* (1x10 minute run), crosshair fixation (1x10 minute run), and movie *The Present* (1x3 minute run) (Alexander et al., 2017). Participants were excluded from analyses if they failed preprocessing (*n*=82) or had less than 8 minutes of low-motion functional data (FD < 0.2mm and outlier frame removal) for each condition (*n*=829), leaving 466 participants for subsequent analyses.

## QUANTIFICATION AND STATISTICAL ANALYSIS

### MRI Data Preprocessing

The MINT precision and HBN consortia datasets were preprocessed using fMRIPrep version 24.1.0 (Esteban et al., 2019) and XCP-D version 0.10.1 (Mehta et al., 2024). The MINT precision data underwent additional denoising prior to fMRIPrep and XCP-D using NORDIC (NOise reduction with DIstribution Corrected PCA) (Moeller et al., 2021), thermal noise for NORDIC was estimated from three noise volumes collected at the end of each functional run, which were removed prior to further preprocessing. As the HBN protocol did not include either phase contrast or noise frames in the functional acquisition as recommended for NORDIC this denoising step was not applied to the HBN consortia data.

### Preprocessing with fMRIPrep

The MINT precision (NORDIC-denoised) and HBN consortia datasets were preprocessed with fMRIPrep version 24.1.0 (Esteban et al., 2019) which is based on Nipype 1.8.6 (RRID:SCR_002502, (Gorgolewski et al., 2011)). A B0-nonuniformity map (or fieldmap) was estimated based on two (or more) spin echo-planar imaging (EPI) references using FSL topup (Andersson et al., 2003).

#### Anatomical data

A single T1-weighted (T1w) image was used per-participant for surface registration. In cases where multiple T1w images were available the ‘best’ image (identified by visual inspection) was used for further preprocessing. The T1w image was corrected for intensity non-uniformity (INU) with *N4BiasFieldCorrection* (Tustison et al., 2010), distributed with *ANTs 2.5.3* (RRID:SCR_004757, (B. Avants et al., 2008)), and used as the T1w-reference throughout the workflow. The T1w-reference was then skull-stripped with a *Nipype* implementation of the *antsBrainExtraction.sh* workflow (from ANTs) using OASIS30ANTs as the target template. Brain tissue segmentation of cerebrospinal fluid (CSF), white-matter (WM), and gray-matter (GM) was performed on the brain-extracted T1w scan using *fast* (FSL (version unknown), RRID:SCR_002823, (Zhang et al., 2001)). Brain surfaces were reconstructed using *recon-all* (FreeSurfer 7.3.2, RRID:SCR_001847, (Dale et al., 1999)), and the brain mask estimated previously was refined with a custom variation of the method to reconcile ANTs-derived and FreeSurfer-derived segmentations of the cortical gray-matter of Mindboggle (RRID:SCR_002438, (Klein et al., 2017)). A T2-weighted image was used to improve pial surface refinement when available. Brain surfaces were reconstructed using recon-all (FreeSurfer 7.3.2, RRID:SCR_001847, (Dale et al., 1999)), and the brain mask estimated previously was refined with a custom variation of the method to reconcile ANTs-derived and FreeSurfer-derived segmentations of the cortical gray matter of Mindboggle (RRID:SCR_002438, (Klein et al., 2017)). Volume-based spatial normalization to two standard spaces (MNI152NLin2009cAsym, MNI152NLin6Asym) was performed through nonlinear registration with antsRegistration (ANTs 2.5.3) using brain-extracted versions of both the T1w reference and the T1w template. The following templates were selected for spatial normalization and accessed with *TemplateFlow* 24.2.0 (Ciric et al., 2022): *ICBM 152 Nonlinear Asymmetrical template version 2009c* [(Fonov et al., 2011), RRID:SCR_008796; TemplateFlow ID: MNI152NLin2009cAsym], *FSL’s MNI ICBM 152 non-linear 6th Generation Asymmetric Average Brain Stereotaxic Registration Model* [(Evans et al., 2012), RRID:SCR_002823; TemplateFlow ID: MNI152NLin6Asym]. *Grayordinate* ‘dscalar’ files containing 91k samples were resampled onto fsLR using Connectome Workbench (Glasser et al., 2013).

#### Functional data

For each BOLD run found per subject (across all tasks and sessions), the following preprocessing was performed. First, a reference volume was identified using a custom methodology of *fMRIPrep* for use in head motion correction. For the multi-echo MINT precision data, the reference volume was from the shortest echo of the BOLD run. Head-motion parameters with respect to the BOLD reference (transformation matrices, and six corresponding rotation and translation parameters) are estimated before any spatiotemporal filtering using *mcflirt* (FSL, (Jenkinson et al., 2002)). The estimated *fieldmap* was then aligned with rigid-registration to the target EPI (echo-planar imaging) reference run. The field coefficients were mapped on to the reference EPI using the transform. The BOLD reference was then co-registered to the T1w reference using *bbregister* (FreeSurfer) which implements boundary-based registration (Greve & Fischl, 2009). Co-registration was configured with six degrees of freedom. The aligned T2w image was used for initial co-registration. This aligned T2* image was carried forward as the *preprocessed BOLD* for the HBN consortia data. Generation of the *preprocessed BOLD* for the multi-echo MINT precision data required an additional multi-echo optimal combination step. Multi-echo optimal combination for the MINT precision data was performed by deriving a T2* map estimated from the preprocessed EPI echoes using tedana’s t2smap workflow (DuPre et al., 2021), by voxel-wise fitting the maximal number of echoes with reliable signal in that voxel to a monoexponential signal decay model with nonlinear regression. The T2*/S0 estimates from a log-linear regression fit were used for initial values. The calculated T2* map was then used to optimally combine preprocessed BOLD across echoes following the method described in (Posse et al., 1999). The optimally combined time series was then carried forward as the *preprocessed BOLD* for the MINT precision data.

Several confounding time-series were calculated based on the *preprocessed BOLD* data: framewise displacement (FD), DVARS, and three region-wise global signals. FD was computed using two formulations following Power and colleagues (absolute sum of relative motions, (Power et al., 2014)) and Jenkinson and colleagues (relative root mean square displacement between affines, (Jenkinson et al., 2002)). FD and DVARS are calculated for each functional run, both using their implementations in *Nipype* (following the definitions by Power et al. 2014). The three global signals are extracted within the CSF, the WM, and the whole-brain masks. Additionally, a set of physiological regressors were extracted to allow for component-based noise correction (*CompCor*, (Behzadi et al., 2007)). Principal components are estimated after high-pass filtering the preprocessed BOLD time-series (using a discrete cosine filter with 128s cut-off) for the two *CompCor* variants: temporal (tCompCor) and anatomical (aCompCor). tCompCor components are then calculated from the top 2% variable voxels within the brain mask. For aCompCor, three probabilistic masks (CSF, WM and combined CSF+WM) are generated in anatomical space. The implementation differs from that of Behzadi et al. in that instead of eroding the masks by 2 pixels on BOLD space, a mask of pixels that likely contain a volume fraction of GM is subtracted from the aCompCor masks. This mask is obtained by dilating a GM mask extracted from the FreeSurfer’s *aseg* segmentation, and it ensures components are not extracted from voxels containing a minimal fraction of GM. Finally, these masks are resampled into BOLD space and binarized by thresholding at 0.99 (as in the original implementation). Components are also calculated separately within the WM and CSF masks. For each CompCor decomposition, the *k* components with the largest singular values are retained, such that the retained components’ time series are sufficient to explain 50 percent of variance across the nuisance mask (CSF, WM, combined, or temporal). The remaining components are dropped from consideration. The head-motion estimates calculated in the correction step were also placed within the corresponding confounds file. The confound time series derived from head motion estimates and global signals were expanded with the inclusion of temporal derivatives and quadratic terms for each (Satterthwaite et al., 2013). Frames that exceeded a threshold of 0.5 mm FD or 1.5 standardized DVARS were annotated as motion outliers. Additional nuisance timeseries are calculated by means of principal components analysis of the signal found within a thin band (crown) of voxels around the edge of the brain, as proposed by (Patriat et al., 2017). The BOLD time-series were resampled onto the left/right-symmetric template “fsLR” using Connectome Workbench (Glasser et al., 2013). *Grayordinates* files (Glasser et al., 2013) containing 91k samples were also generated with surface data transformed directly to fsLR space and subcortical data transformed to 2 mm^3^ resolution MNI152NLin6Asym space. All resamplings can be performed with a *single interpolation step* by composing all the pertinent transformations (i.e. head-motion transform matrices, susceptibility distortion correction when available, and co-registrations to anatomical and output spaces). Gridded (volumetric) resamplings were performed using *nitransforms*, configured with cubic B-spline interpolation. Many internal operations of fMRIPrep use Nilearn 0.10.4 (RRID:SCR_001362, (Abraham et al., 2014)), mostly within the functional processing workflow.

#### Post-processing of fMRIPrep Outputs with XCP-D

The MINT precision and HBN consortia datasets then underwent further post-processing with the eXtensible Connectivity Pipeline-DCAN (XCP-D) version 0.10.1 (Mehta et al., 2024; Ciric et al., 2018; Satterthwaite et al., 2013). XCP-D was built with Nipype version 1.8.6 ((Gorgolewski et al., 2011), RRID:SCR_002502).

### Anatomical data

Native-space T1w images were transformed to MNI152NLin2009cAsym space at 1 mm^3^ resolution. fsLR-space morphometry surfaces were copied from the preprocessing derivatives to the XCP-D derivatives. HCP-style midthickness, inflated, and very-inflated surfaces were generated from the white-matter and pial surface meshes. fsnative-space surfaces were then warped to fsLR space.

### Functional data

For each of the BOLD runs found per subject (across all tasks and sessions), the following post-processing was performed. The six translation and rotation head motion traces were band-stop filtered to remove signals between 12.0 and 16.0 (MINT precision data, automatically modified from 12.0 and 18.0 BPM due to Nyquist frequency constraints) or 12.0 and 18.0 (Healthy Brain Network consortia data) breaths-per-minute using a(n) first-order notch filter, based on (Fair et al., 2020). The Volterra expansion of these filtered motion parameters was then calculated. Framewise displacement was calculated from the filtered motion parameters using the formula from (Power et al., 2014) with a head radius of 50 mm. Volumes with filtered framewise displacement greater than 0.2 mm were flagged as high-motion outliers for the sake of later censoring (Power et al., 2014). In total, 36 nuisance regressors were selected from the preprocessing confounds, according to the ‘36P’ strategy. These nuisance regressors included six motion parameters, mean global signal, mean white matter signal, mean cerebrospinal fluid signal with their temporal derivatives, and quadratic expansion of six motion parameters, tissue signals and their temporal derivatives (Ciric et al., 2017; Satterthwaite et al., 2013). Any motion parameters in the confounds file were filtered using the same parameters as described above and the Volterra expansion was calculated. The BOLD data were converted to NIfTI format, despiked with *AFNI’s 3dDespike*, and converted back to CIFTI format. Nuisance regressors were regressed from the BOLD data using a denoising method based on *Nilearn’s* approach. Any volumes censored earlier in the workflow were first cubic spline interpolated in the BOLD data. Outlier volumes at the beginning or end of the time series were replaced with the closest low-motion volume’s values, as cubic spline interpolation can produce extreme extrapolations. The timeseries were band-pass filtered using a(n) second-order Butterworth filter, in order to retain signals between 0.009-0.08 Hz. The same filter was applied to the confounds. The resulting time series were then denoised via linear regression, in which the low-motion volumes from the BOLD time series and confounds were used to calculate parameter estimates, and then the interpolated time series were denoised using the low-motion parameter estimates. Postprocessing derivatives from multi-run tasks were then concatenated across runs and directions. Many internal operations of XCP-D use AFNI (Cox, 1996; Cox & Hyde, 1997), Connectome Workbench (Glasser et al., 2013), ANTS (B. Avants et al., 2008; B. B. Avants et al., 2009), TemplateFlow version 24.2.2 (Ciric et al., 2022), matplotlib version 3.9.2 (Hunter, 2007), Nibabel version 5.3.0 (Brett et al., 2024), Nilearn version 0.10.4 (Abraham et al., 2014), numpy version 2.1.2 (Harris et al., 2020), pybids version 0.17.2 (Yarkoni et al., 2019), and scipy version 1.14.1 (Virtanen et al., 2020). For more details, see the XCP-D website (https://xcp-d.readthedocs.io).

### Individual Large-Scale Functional Network Topography Estimation with ReproTM

Individual large-scale functional networks were generated for each participant by viewing condition using the template matching network detection algorithm (Gordon, Laumann, Adeyemo, & Petersen, 2017) from the Reproducible Template Matching (ReproTM) Toolbox version 1.0.0 (Godfrey, 2026). The input for template matching is dense connectivity matrix (here, 59k x 59k cortical vertices) which was generated per-participant and per-viewing condition using input dense time series data that underwent surface smoothing (FWHM = 2mm) and censoring. The censoring mask was generated by assessing temporally filtered framewise displacement (FD) values to identify high motion frames (FD > 0.2mm) and outlier frames (FD > 3 scaled median absolute deviation) which were scrubbed from the timeseries. Dense time series data were then matched for length (total retained frames) within participants across viewing conditions by truncating the longer timeseries. Matching total retained frames of the input dense time series was intended to partially account for effects of scan duration on functional connectivity reliability across viewing conditions. Finally, prior to running template matching, the input dense connectivity matrix was z-scored separately for each hemisphere, the subcortex, and connections between the cortex and subcortex to account for potential drop out and reduced subcortical signal as previously described (Hermosillo et al., 2024).

The ReproTM Toolbox version 1.0.0 (Godfrey, 2026) was then used to match each participant’s cortical dense connectivity to a set of network templates with the template matching network detection algorithm (Gordon, Laumann, Adeyemo, & Petersen, 2017). In brief, template matching calculates the similarity between seedmaps of each vertex to the seedmap of every template network (here, template seedmaps were thresholded to include only the top cortical connections *Z* > 1.0). Vertex-to-network similarities were calculated using sum of squares variance decomposition to produce an intermediate matrix with similarities (η^2^) for each vertex to every template network. Network assignments were then determined with a winner-take-all procedure, where each vertex was assigned to the template network with the maximal similarity value. The template networks used in the present study were generated using independent data from the ABCD consortia (n = 141, mean age = 9.92 ± 0.63 years; 67 female/ 74 male) in prior work (Hermosillo et al., 2024). Template networks included the default mode (DMN), visual (Vis), frontoparietal (FP), dorsal attention (DAN), language (Lang), salience (Sal), action mode (AMN), somatomotor dorsal (SMd), somatomotor lateral (SMl), auditory (Aud), temporal pole (Tpole), medial temporal lobe (MTL), parietal memory (PMN), parietal occipital (PON), and somato-cognitive action (SCAN). A graphical representation of the template matching method used herein is provided in the supplemental materials (Figure S3).

### Analysis Overview

#### Individualization and Reliability (Figure 2)

We assessed individualization (i.e., increase in same individual similarity over similarity to other individuals) and reliability of large-scale functional networks between movie watching and crosshair fixation in the MINT precision data by first calculating similarity values between all possible observation pairs (19 participants x 2 viewing conditions x 2 sessions). Functional network similarities were compared at the levels of network assignments (network assignment vectors compared with Dice correlation), vertex-to-network similarity matrices (vectorized upper triangle compared with linear bivariate correlation), and dense connectivity matrices (vectorized upper triangle compared with linear bivariate correlation) to create a second order ‘similarity matrix’ (Figure 2A) following a prior application of this method for functional connectivity (Gordon, Laumann, Gilmore, et al., 2017; Gratton et al., 2018).

Individualization for both movie watching and crosshair fixation was determined using paired sample t-tests to compare same individual similarities (per-participant test-rest reliability across data sessions, by condition) to the mean different individual similarities (per-participant mean correlation to all other individuals, by condition). Reliability between movie watching and crosshair fixation was tested using a paired sample t-test to compare the same individual similarities (per-participant test-retest reliability across data sessions, by condition) between movie watching and crosshair fixation (Figure 2B).

#### Border and Ectopic Variants (Figure 4)

Variations in large-scale functional network estimates within-participants during movie watching were characterized as either border shifts or ectopic intrusions relative to participant’s own crosshair fixation baseline by adapting publicly available code (https://github.com/GrattonLab/Dworetsky_BorderEctopicVariants) that was developed to characterize between-participant differences relative to a group average baseline (for details, see (Dworetsky et al., 2024)). In brief, within each participant, vertices which varied in network assignment between movie watching and crosshair fixation were binarized (variant or non-variant) and grouped into contiguous clusters of at least 30 vertices that shared the same network assignment in the movie watching condition. If any vertex within a cluster shared the same network assignment with a neighboring vertex, or with a vertex within 3.5mm, that cluster was labeled as a border shift. Clusters which did not meet the above criteria were labeled as ectopic intrusions.

#### Cortical Network Surface Area (Figure 5, Figure 8B)

Cortical network surface areas were derived per-participant and per-viewing condition for each of the 15 large-scale functional networks considered. The surface area of each individual vertex in mm^2^ was derived from participant’s own midthickness files using the connectome workbench -*surface-vertex-areas* function. The assignment of vertices to one of the 15 functional networks was derived using the ReproTM toolbox as described above. Cortical surface areas for each network were then derived by summing surface area values across all vertices assigned to the same network. Cortical surface area values for each network were then transformed to a proportion of the total cortical surface area (sum of surface area values across all vertices regardless of network assignment) to account for differences in total cortical surface area across participants.

The effect of viewing condition on cortical network surface area was examined using separate linear mixed effects models for each network, with cortical surface area as the outcome and a main effect of viewing condition (movie watching > crosshair fixation) while covarying for fixed effects of age, sex, and head motion (mean framewise displacement in mm) and random subject intercepts. Probability values were false discovery rate corrected across linear mixed effects models (15 total models, one per network). These analyses were repeated within each MINT independent data session and in the HBN consortia data using the same main effect and covariates.

#### Whole Cortex Magnitude of Group, Individual, and Viewing Condition Effects (Figure 6A, Figure 8C)

The magnitude of group, individual, and viewing condition ‘effects’ across the whole cortex were derived from the network assignment similarity matrix (Figure 2A) following an approach previously developed and applied to functional connectivity data (Gratton et al., 2018). In the MINT precision data, similarity between whole cortex network assignment vectors was calculated using Dice correlation across observations paired to estimate the following effects: 1) group effect (mean similarity of networks from different individuals, sessions, and conditions), 2) condition effect (mean similarity of networks from different individuals and sessions but the same condition), 3) individual effect (mean similarity of networks from the same individual but different conditions), 4) individual by session effect (mean similarity of networks from the same individual and session but different conditions), and 5) individual by condition effect (mean similarity of networks from the same individual and condition) (Figure 6A). The HBN consortia data were used to estimate group, condition, and individual effects using a similar approach (Figure 8C); however, we note that the HBN data is collected at a single session and therefore lacks the test retest design necessary to derive the individual by condition effect (i.e., increased same individual similarity for functional network topography derived from the same viewing condition across data sessions) so the individual by condition effect could not be tested for replicability.

#### Whole Cortex Normalized Relative Effect Magnitude of Group, Individual, and Viewing Condition Effects (Figure 6B, Figure 8C)

The magnitude of group, individual, and viewing condition ‘effects’ across the whole cortex were contrasted to identify the normalized relative magnitude of each effect, paralleling prior work with connectivity (Gratton et al., 2018). The relative magnitude of each effect was first calculated by comparing the improvement in similarity values of each effect to its baseline (red lines in Figure 6A, Figure 8). In the MINT precision data, the group effect served as the “baseline” for the individual and condition effects (i.e., individual and condition effects were estimated as added similarity above the group effect), while the individual effect served as the “baseline” for the individual by session and individual by condition effects (i.e., individual by session and individual by condition effects were estimated as added similarity above the individual effect). These relative effects were then normalized as a proportion of the maximum similarity (for the MINT precision data, the individual by condition similarity) to calculate a normalized relative effect magnitude (Figure 6B). In the HBN consortia data, the group effect served as the “baseline” for the individual and condition effects and these relative effects were normalized as a proportion of the maximum similarity (for the HBN consortia data, the individual similarity) (Figure 8C). The HBN consortia data lacked the test-retest design necessary to estimate individual by session and individual by condition effects, so these normalized relative effect magnitudes were not derived.

#### Variable Cortical Spatial Distributions of Group, Individual, and Viewing Condition Effects (Figure 6C, Figure 8C)

To examine how group, individual, and viewing condition effects varied across the cortical surface, we used the same approach described above for deriving normalized relative effect magnitudes (Gratton et al., 2018) at each individual vertex. In the MINT precision data, at each vertex, the same approach of calculating the average paired similarity was applied to the vertex-to-network similarities (1 x 15 vector; representing similarity values (η^2^) for each vertex to all template networks) (see Figure S3). The similarity of vertex-wise continuous vertex-to-network similarity vectors was calculated across observations (19 participants x 2 sessions x 2 conditions) using linear bivariate correlations paired as described above to estimate normalized relative effects of group, condition, individual, individual by session, and individual by condition relative to the same effect “baselines” and normalized as a proportion for the maximum individual by condition similarity. In the HBN consortia data, vertex-wise continuous vertex-to-network similarity vectors were also examined using a paired similarity approach to calculate normalized relative effect magnitudes of group, condition, and individual (following “baselining” of the condition and individual similarity above the group similarity) which were normalized relative to the maximum individual by condition similarity.

#### Group, Individual, and Viewing Condition Effects in Clinically Relevant Regions of Interest (Figure 7)

To test whether the contribution of group, individual, and viewing condition effects varied within clinically relevant regions of interest we used independent data from *Neurosynth* (Yarkoni et al., 2011) to generate masks for three common neuromodulation targets: somatomotor cortex (search term: “sensorimotor cortex”; test of voxel activation: z-scored association test; probability value = 0.01, FDR corrected), dorsolateral prefrontal cortex (search term: “dorsolateral pfc”; test of voxel activation: z-scored association test; probability value = 0.01, FDR corrected), and temporal parietal junction (search term: “temporoparietal junction”; test of voxel activation: z-scored association test; probability value = 0.01, FDR corrected). Clinically relevant regions of interest masks were transformed from volume to surface space using in-house scripts to call functions from Connectome Workbench (Glasser et al., 2013). The volume template was first transformed into left and right surface files using the *-volume-to-surface-mapping* function after which the left and right surface files were combined using the *-cifti-create-dense-from-template* function to create a surface mask for each region (Figure 7A). We then applied the same approach for estimating whole cortex normalized relative effect magnitudes by masking the whole cortex network assignment vectors to only include vertices within each clinically relevant region of interest which were examined separately (Figure 7B-D)

## Supporting information

Full Supplement

