## Supplementary material for "Movie Watching for Individual Functional Brain Network Estimation in Adolescents": Full Supplement

### SUPPLEMENTAL FIGURES AND TABLES

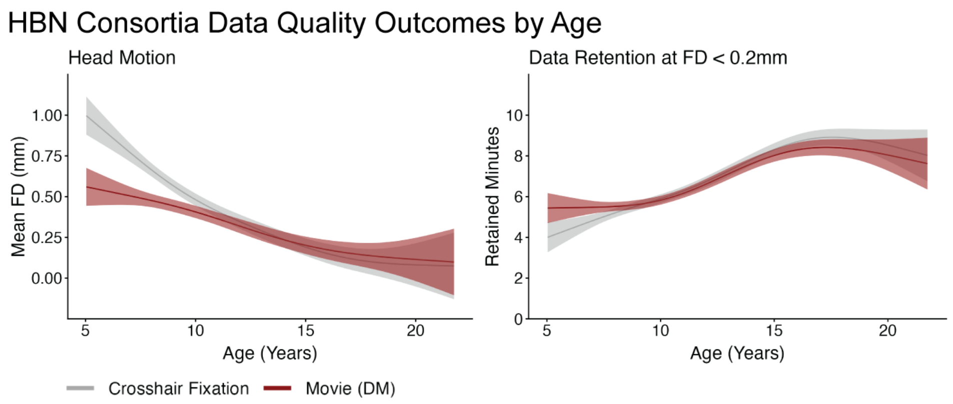


**Figure S1 (Related to Figure 1). Movie Watching Effects on Functional Data Quality Outcomes by Age, HBN Consortia Data.** Examination of non-linear viewing condition effects on functional data quality outcomes including head motion (mean framewise displacement, mm) and functional data retention (minutes retained at FD < 0.2mm) suggest that movies have greater benefit for improving functional data quality in younger children replicating prior reports (Greene et al., 2018; Frew et al., 2022).

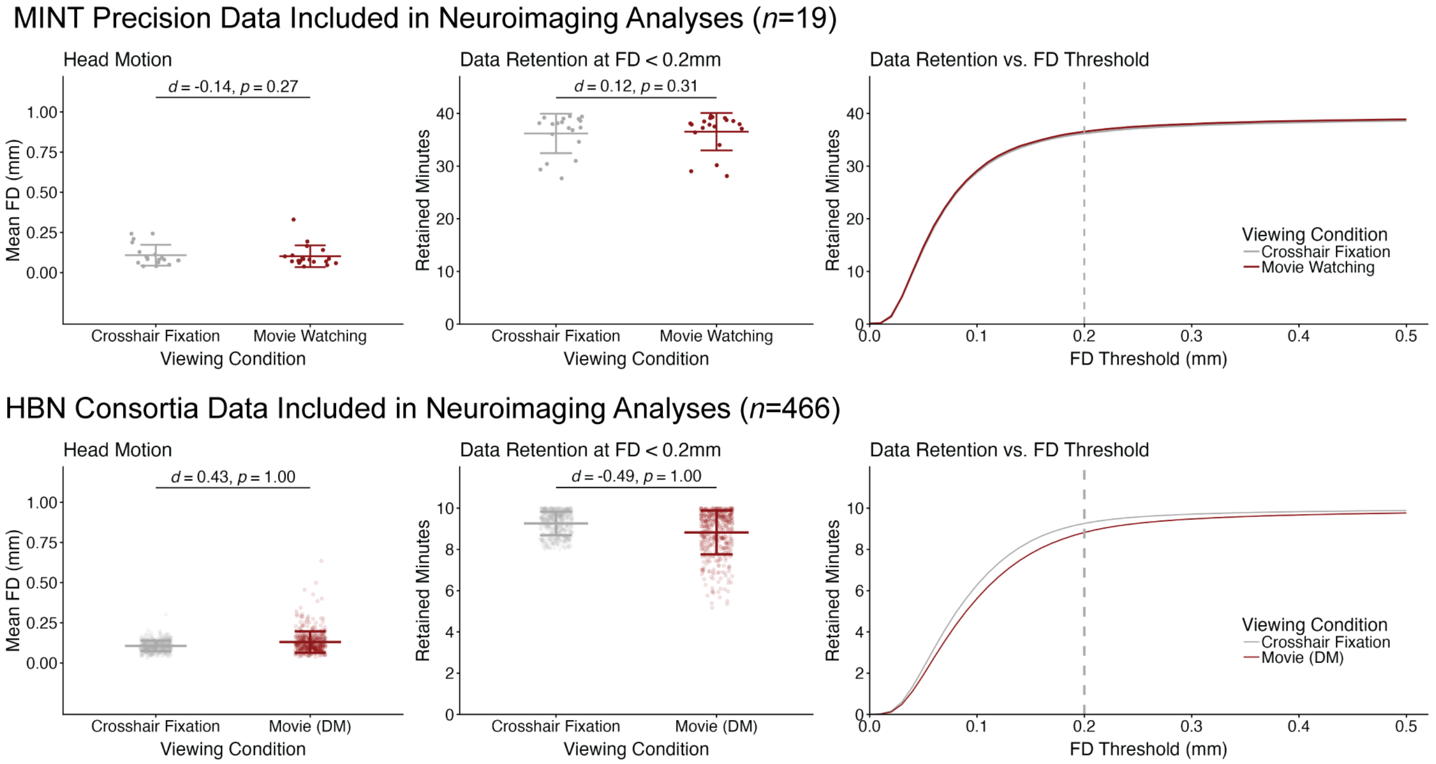


**Figure S2 (Related to Figure 1). Similar Functional Data Quality Across Movie Watching and Crosshair Fixation for Participants Included in Neuroimaging Analyses.** There were no effects of viewing condition on functional data quality after data quality exclusions (excluded participants; MINT precision data *n*=9, HBN consortia data *n*=812) for head motion (mean framewise displacement in mm, meanFD) in the MINT precision data (meanFD; crosshair fixation mean = 0.11, movie clips mean = 0.10) and HBN consortia data (meanFD; crosshair fixation mean = 0.11, movie DM mean = 0.13). There were additionally no effects of viewing condition on functional data retained at the study threshold of FD < 0.2mm (post-censored minutes) in the MINT precision (post-censored minutes; crosshair fixation mean = 36.19, movie watching mean = 36.54) and HBN consortia data (post-censored minutes, crosshair fixation mean = 9.26, movie DM mean = 8.82). We note that although minutes of post-censored functional data are quantified here as a measure of data quality, functional data were matched for length within participants across viewing conditions prior to further neuroimaging analyses.

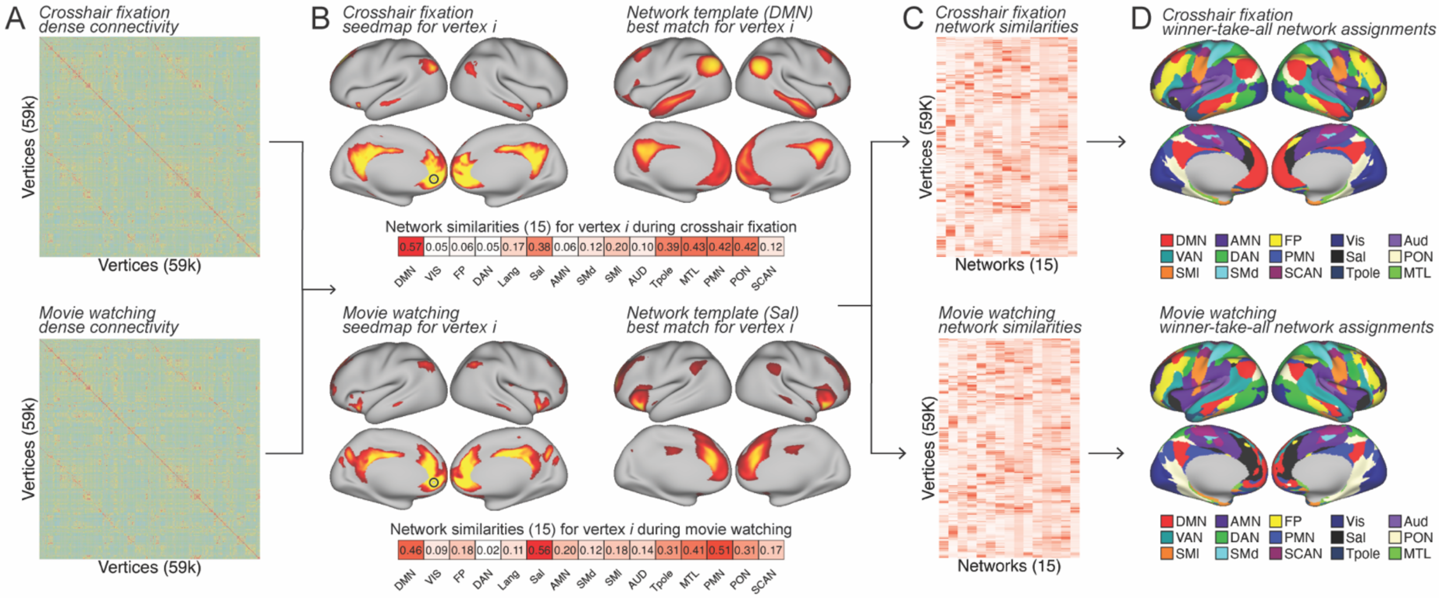


**Figure S3. Individual Large-scale Functional Network Topography Estimation with Template Matching.** A) Cortical dense functional connectivity was derived from functional MRI collected for crosshair fixation (white cross, black background) or movie watching (concatenated *Pixar* shorts for children). B) Spatial similarities (here, $\eta^2$ derived with sum of squares variance decomposition) were calculated between all vertex seedmaps (visualized for exemplar vertex *i*) to each network template (visualized for exemplar vertex *i*'s best network matches). C) The template matching procedure produces an intermediate similarity matrix for each vertex to every template network. D) Network assignments are determined with a winner-take-all heuristic where each vertex is assigned to the template network with the greatest $\eta^2$ similarity.

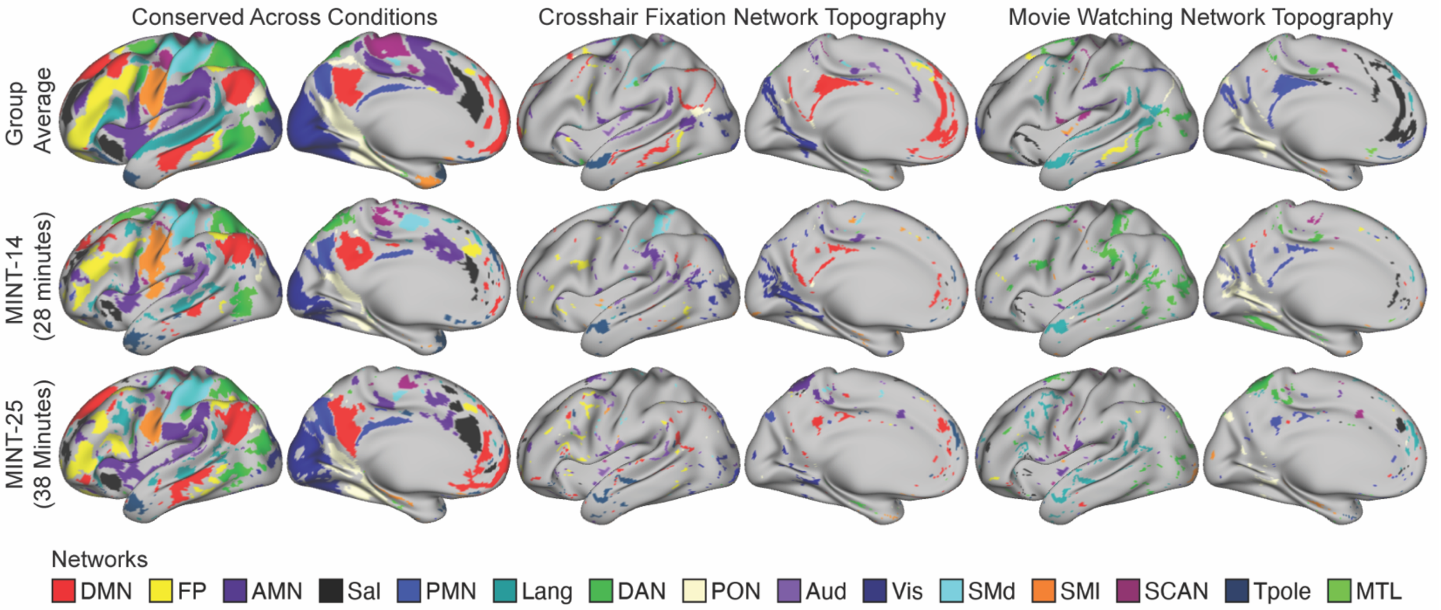


**Figure S4. Strict Reproducibility of Functional Network Topography Across Crosshair Fixation and Movie Watching, MINT Precision Data.** A strict vertex-wise reproducibility criteria was applied to characterize highly reproducible patterns of functional network topography across viewing conditions. Individual vertices were considered reproducibly conserved across conditions if they had the same functional network assignment in both viewing conditions at both independent data sessions (e.g., vertex *i* was assigned to default mode network for crosshair fixation session 1, crosshair fixation session 2, movie watching session 1 and movie watching session 2). Individual vertices were considered reproducibly reassigned if they had the same functional network reassignment at both independent data sessions (e.g., vertex *i* was assigned to default mode network in crosshair fixation session 1 and crosshair fixation session 2; and the same vertex *i* was reassigned to language network in movie watching session 1 and movie watching session 2). Data are visualized for the group average and the same exemplar participants with varying degrees of low-motion data at each independent data session (MINT-14: 14 minutes per-condition at each data session; MINT=25: 19 minutes per-condition at each data session).

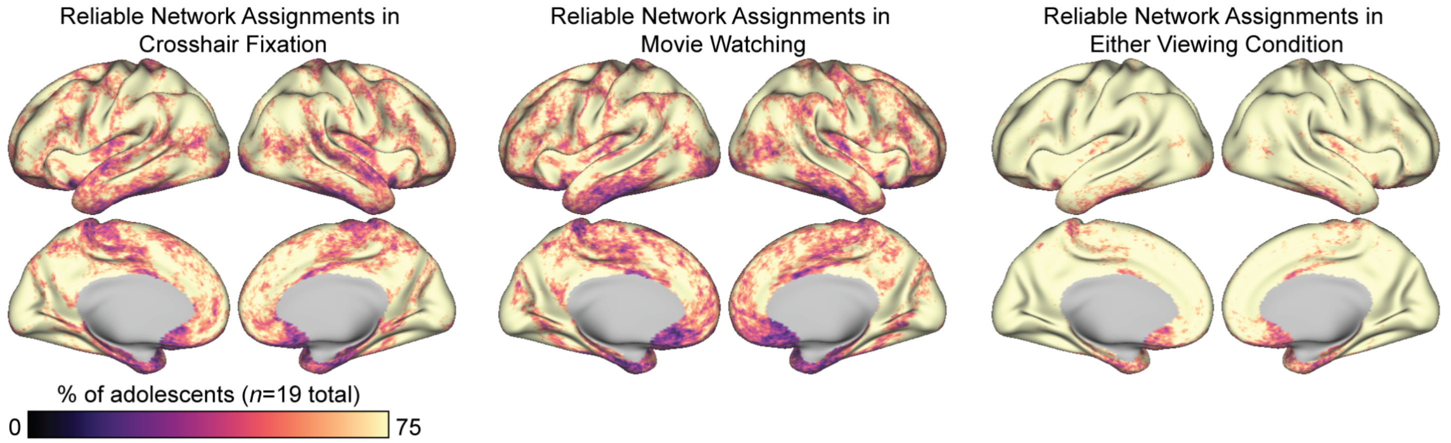


**Figure S5. (Related to Figure 2). Functional Network Topography Reliability at Individual Vertices for Movie Watching and Crosshair Fixation, MINT Precision Data.** Density maps showing the proportion of adolescents (% of total) with reliable functional network topography (winner-take-all network assignments) at individual vertices. At individual vertices, for each adolescent independently by viewing condition, vertices were determined to have reliable functional network topography if the vertex was assigned to the same winner-take-all functional network assignment at both independent data sessions. These vertex-wise reliability binary maps (1 = reliable, 0 = not reliable) for each viewing condition were averaged across adolescents and converted to a proportion of the total sample. We additionally present a density map showing vertices that were reliable in either crosshair fixation or movie watching calculated using a similar approach.

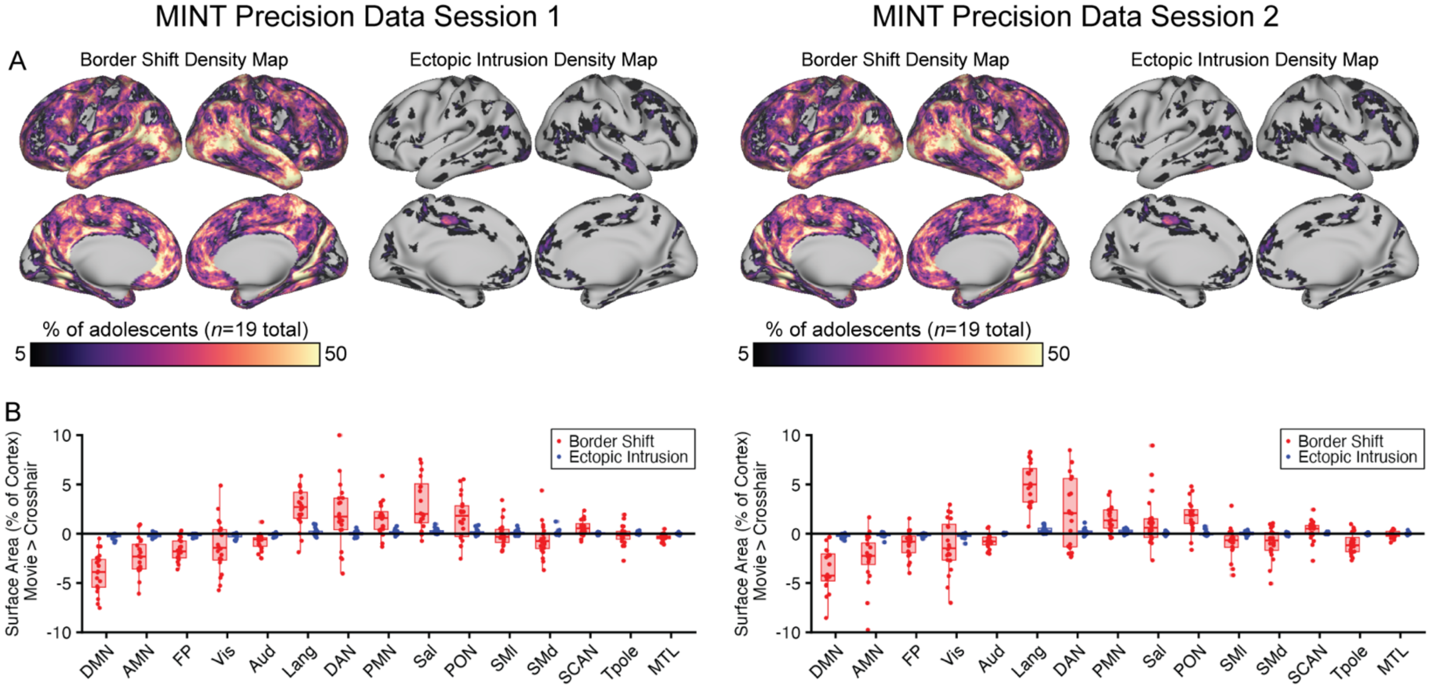


**Figure S6. (Related to Figure 4). Border and Ectopic Variants Are Reproducible Across Independent Sessions, MINT Precision Data.** A) Density maps, showing the proportion of adolescents with a border shift or ectopic intrusion at each individual vertex, demonstrate similar spatial distribution and magnitude of border shifts and ectopic intrusions across independent data sessions. B) Network reassignments between movie watching and crosshair fixation were primarily border shifts (red) over ectopic intrusions (blue) and the magnitude of border shifts and ectopic intrusions were largely comparable across independent data sessions. *DMN:* default mode network, *AMN:* action mode network; *FP*: frontoparietal network; *Aud:* auditory network; *Lang*: language network; *DAN:* dorsal attention network; *PMN:* parietal memory network; *Sal:* salience network; *PON:* parietal occipital network; *SMl:* somatomotor lateral network; *SMd:* somatomotor dorsal network; *SCAN:* somato-cognitive action network; *TPole:* temporal pole; *MTL:* medial temporal lobe.

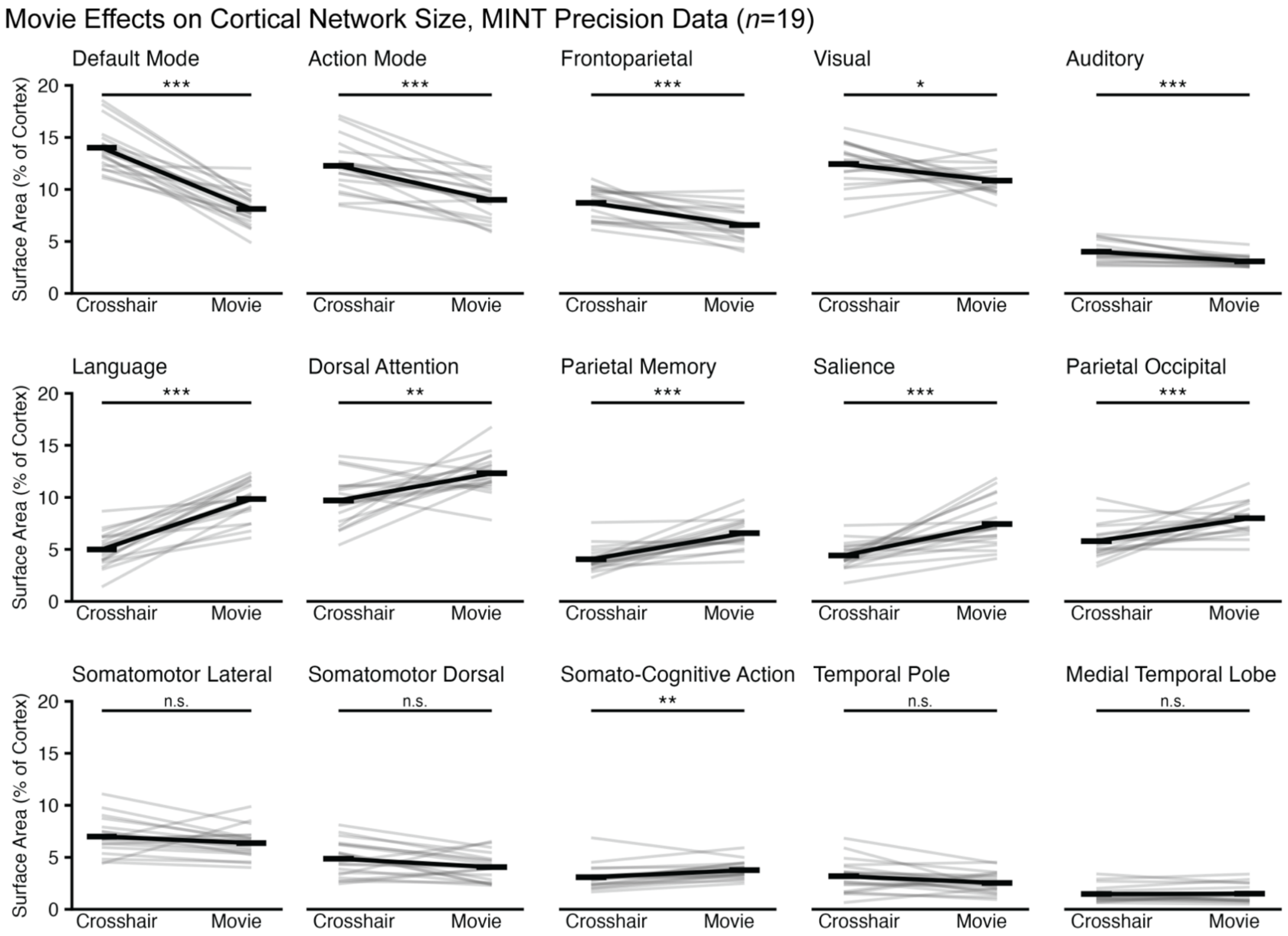


**Figure S7. (Related to Figure 5). Cortical Network Surface Area Across Movie Watching and Crosshair Fixation, MINT Precision Data.** Several movie clip effects on cortical network surface area estimates detected in the group average (black lines) were also highly consistent across individual participants (colored lines). Group average patterns with notably high consistency across individuals (% of total sample) were decreased cortical surface area of default mode 100%), action mode (95%), and frontoparietal (95%) networks and increased cortical surface area of language (100%), parietal memory (95%), and salience (89%) networks. Statistical significance was determined using separate linear mixed effects models for each network, with cortical surface area as the outcome with a main effect of viewing condition while covarying for fixed effects of age, sex and head motion (mean framewise displacement), and random subject intercepts. False discovery rate correction was applied across the 15 network linear mixed effects models. ***: $p_{FDR} < 0.001$; **: $p_{FDR} < 0.01$; *: $p_{FDR} < 0.05$; n.s.: $p_{FDR} > 0.05$.

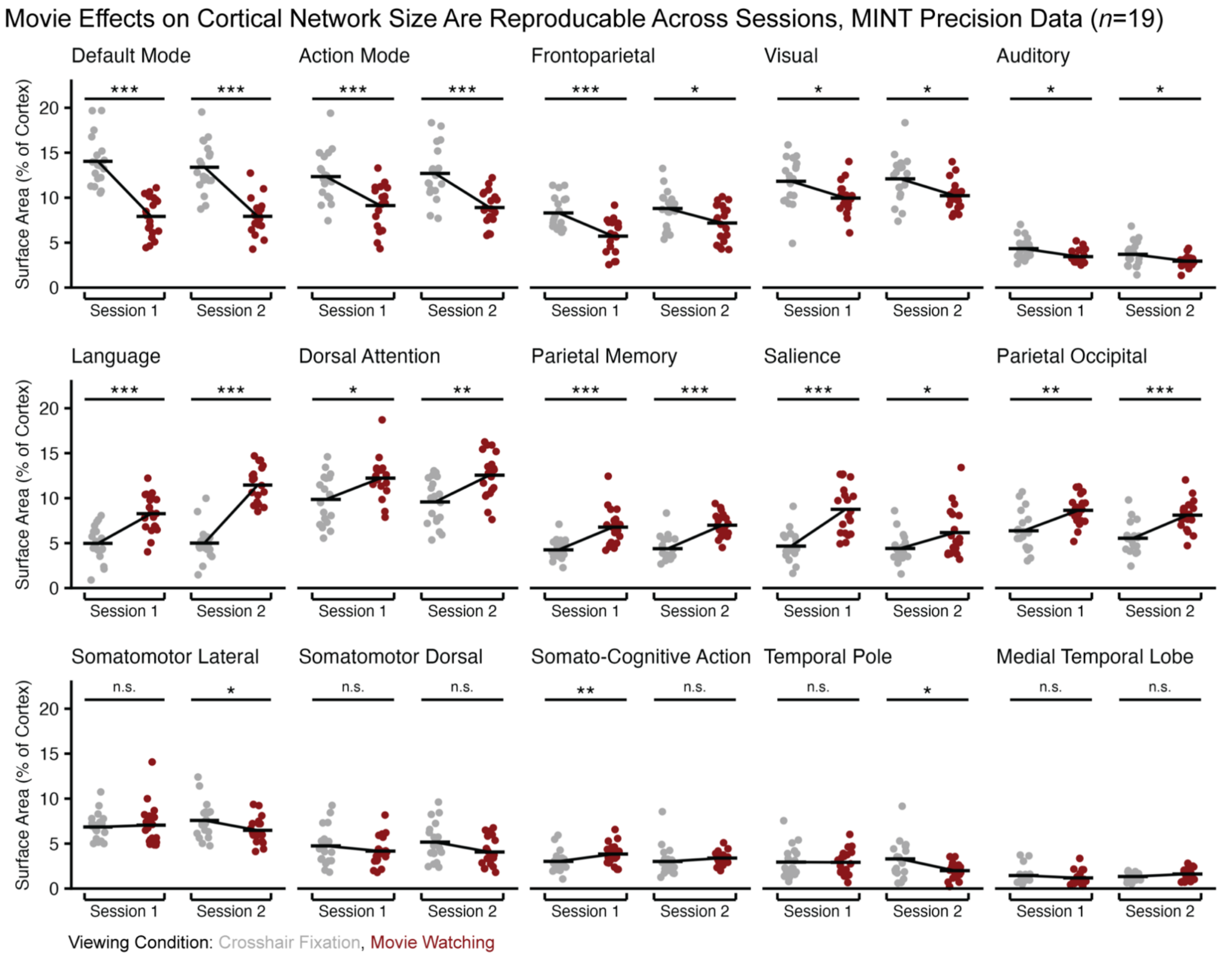


**Figure S8. (Related to Figure 5). Movie Effects on Cortical Network Size Are Reproducible Across Independent Data Sessions, MINT Precision Data**. Data points represent network surface area estimates for individual participants (*n*=19) at each independent data session, black lines represent surface area means across participants per-condition and per-session. Statistical significance was determined using separate linear mixed effects models per network and per session with cortical network surface area as the outcome predicted by viewing condition (movie watching > crosshair fixation) while covarying for fixed effects of age, sex, head motion (mean framewise displacement in mm), and random subject intercepts. False discovery rate correction was applied across the 15 network linear mixed effects models separately for each independent session. ***: $p_{FDR} < 0.001$; **: $p_{FDR} < 0.01$; *: $p_{FDR} < 0.05$; *n.s.*: $p_{FDR} > 0.05$.

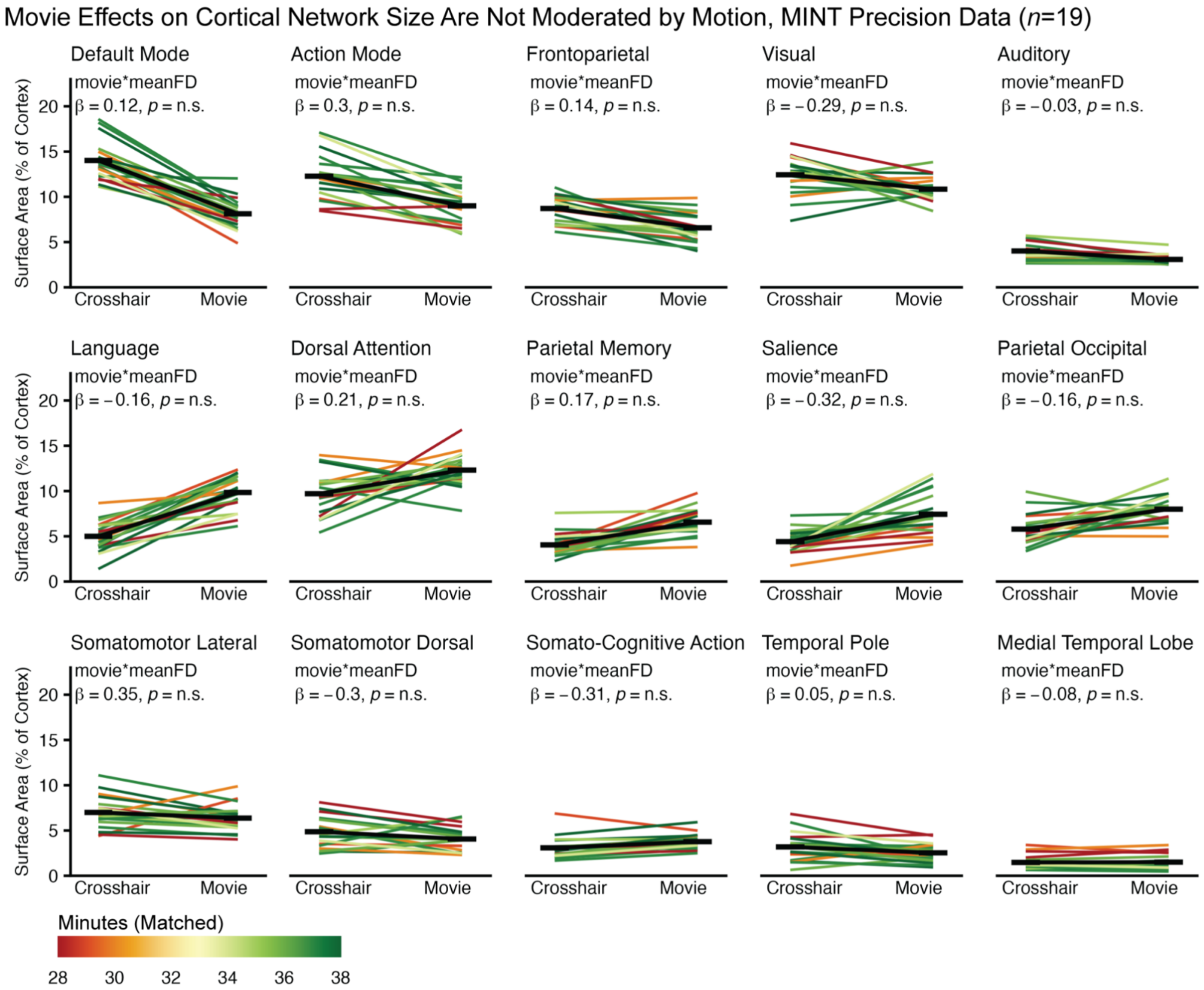


**Figure S9 (Related to Figure 5). No Moderating Effect of Head Motion on the Relation Between Condition and Cortical Network Surface Area, MINT Precision Data**. Cortical network surface area (% of total in $mm^2$) between crosshair fixation and movie watching for each participant in the MINT precision data (*n*=19). Black lines represent across-participant means by viewing condition and colourful lines represent individual participants, coloured by retained post-censored functional data at the study threshold of FD < 0.2mm. Within participants in the MINT precision data, to partially account for motion effects, the dense time series values were matched for length across viewing conditions by truncating the longer time series which resulted in one value for matched minutes per-participant. The interaction between the movie watching and head motion for predicting cortical network surface area was examined with separate linear mixed effects models for each network, with cortical surface area as the outcome with a main effect of condition (movie watching > crosshair fixation), head motion (mean framewise displacement, in mm), the interaction between condition and head motion, while covarying for fixed effects of age and sex and random subject intercepts. False discovery rate correction was applied across the 15 network linear mixed effects models. *meanFD*: mean framewise displacement, n.s.: $p_{FDR} > 0.05$.

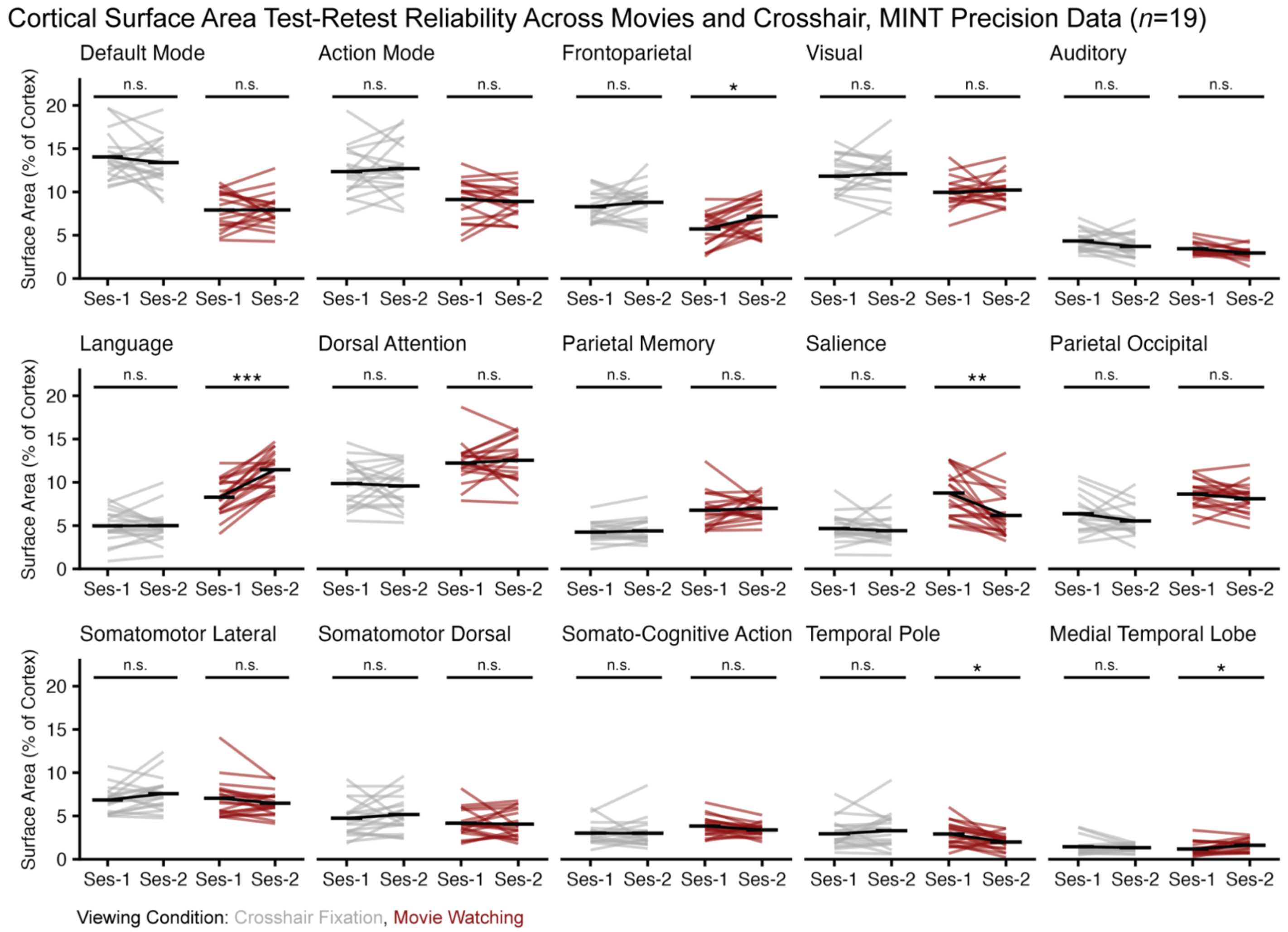


**Figure S10 (Related to Figure 3). Session Effects on Cortical Network Surface Area Across Movie Watching and Crosshair Fixation, MINT Precision Data.** Cortical network surface area (% of total in mm2) were derived for each MINT precision participant (*n*=19) for crosshair fixation and movie watching at each independent data session (13–19 minutes of low-motion data per session, per condition, per participant). Black lines represent across-participant means while colourful lines represent individual participants. The effect of session on cortical network surface area was examined with separate linear mixed effects models per network and condition, with cortical network surface area as the outcome with a main effect of session while covarying for fixed effects of sex, age, and head motion (mean framewise displacement in mm) and random subject intercepts. Probability values for the session effect were false discovery rate corrected across the 15 network linear mixed effects models separately for each viewing condition. Ses-1: Independent data session 1; Ses-2: Independent data session 2; ***: $p_{FDR} < 0.001$; **: $p_{FDR} < 0.01$; *: $p_{FDR} < 0.05$; n.s.: $p_{FDR} > 0.05$.

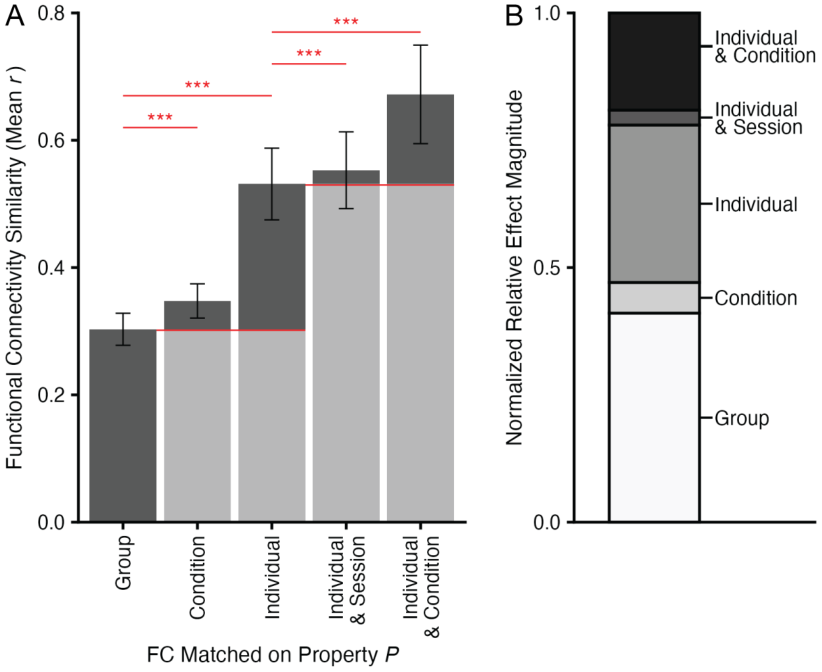


**Figure S11 (Related to Figure 6). Magnitude of Group, Individual, and Condition Effects on Functional Connectivity, MINT Precision Data.** In addition to our analysis on the main study outcome of functional network topography (vertex-wise network assignments) (Figure 6), we also characterized condition effects relative to condition-invariant group and individual effects for explaining variance in functional connectivity (vectorized upper triangle of the dense cortical connectome, 59k by 59k) as in prior work (Gratton et al., 2018). A) Mean similarity of whole-cortex vectorized dense functional connectivity matrices at the level of group (different adolescents, different conditions), condition (different adolescents, same condition), individual (same adolescent, different conditions), individual by session (same adolescent, different conditions, but the same session), and individual by condition (same adolescent, same condition). B) The improved similarity of whole-cortex dense functional connectivity as quantified relative to the maximum similarity (here, the individual by condition similarity) to get normalized relative magnitude of each additive 'effect' following the approach of (Gratton et al., 2018). As with functional network topography, the largest contributors to dense functional connectivity were condition-invariant group and individual effects. However, unlike functional network topography where group effects were much larger than individual effects, when considering dense functional connectivity the relative magnitude of group effects (41% normalized relative variance) and individual effects (31% normalized relative variance) were more similar.

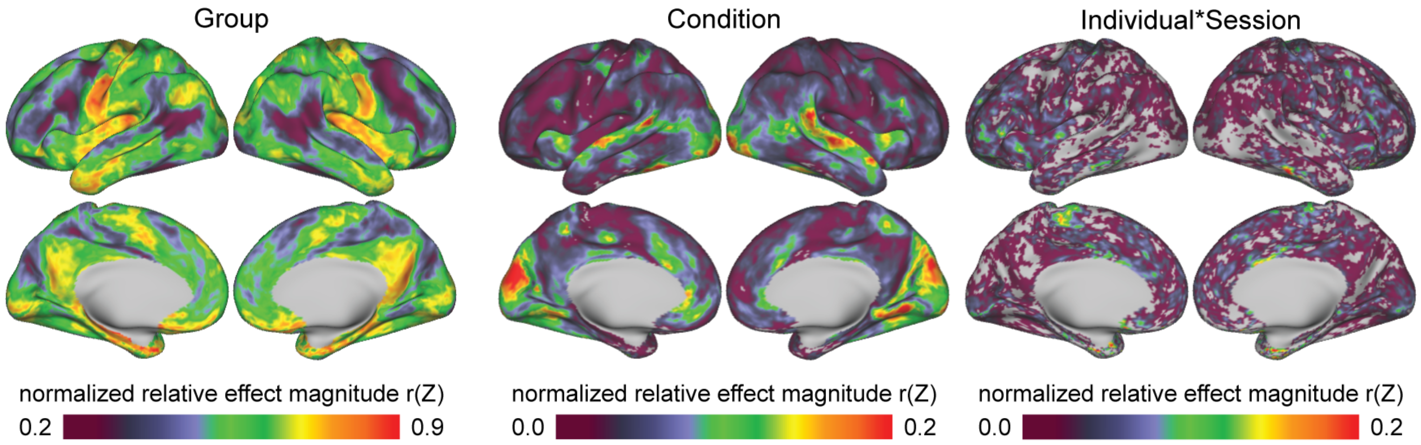


**Figure S12. (Related to Figure 6). Spatial Distributions of Group, Condition, and Individual by Session Effects with Variable Scaling, MINT Precision Data.** The same cortical spatial distributions of normalized relative group and condition effects (Figure 6), alongside individual by session effects, are presented using custom color scales to better visualize each effects spatial distribution across the cortical surface.

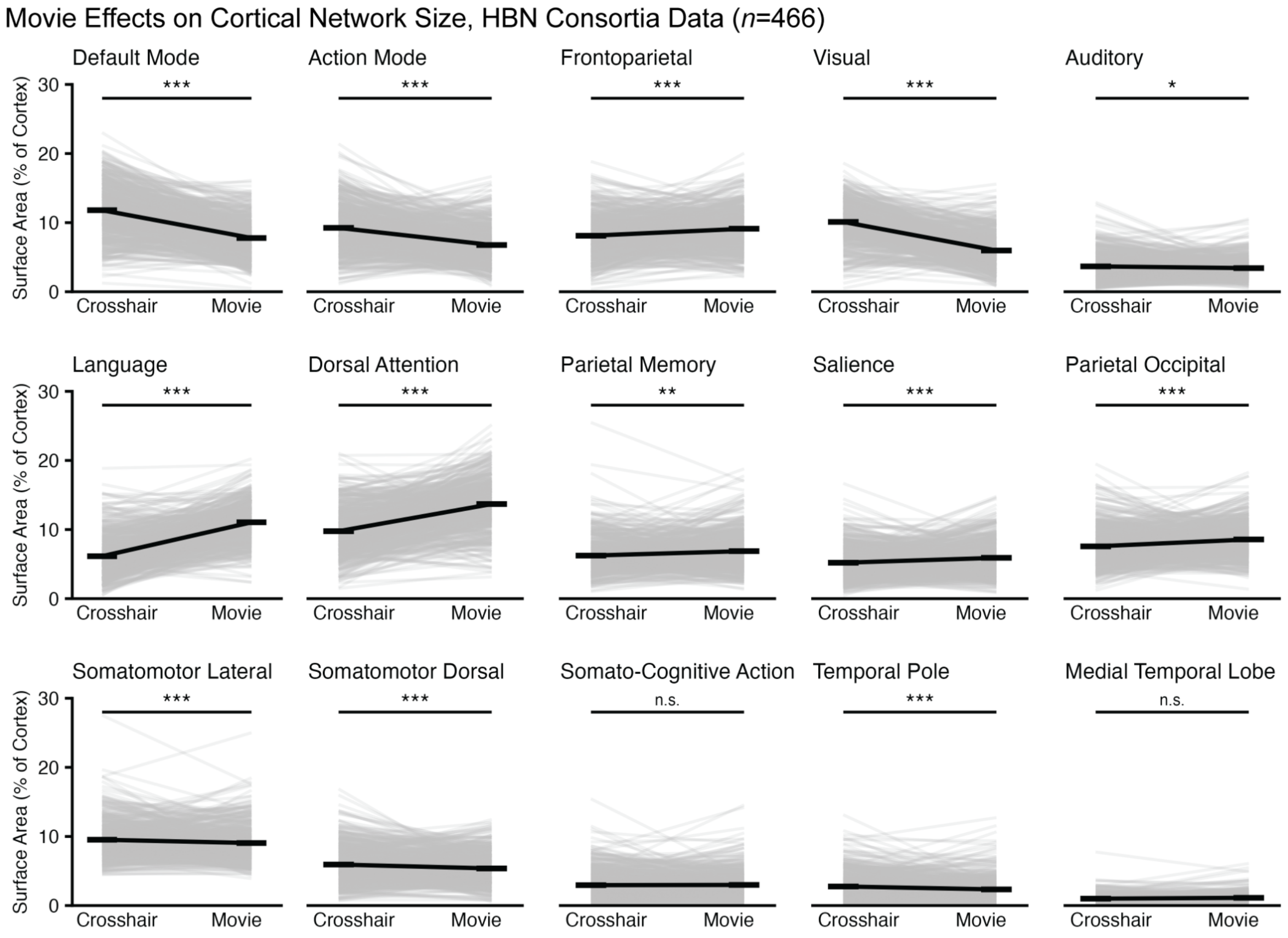


**Figure S13. (Related to Figure 8). Movie Effects on Network Surface Area, Healthy Brain Network Consortia Data.** Cortical network surface area estimates (% of total) in the HBN Consortia Data ($n$ = 466) between crosshair fixation and movie watching are visualized for individual participants (grey lines) and the mean across participants (black lines). Statistical significance was determined using separate linear mixed effects models per network with cortical surface area as the outcome predicted by viewing condition while covarying for fixed effects of age, sex, and head motion (mean framewise displacement), and random subject intercepts. False discovery rate correction was applied across the 15 network linear mixed effects models. ***: $p_{FDR} < 0.001$; **: $p_{FDR} < 0.01$; *: $p_{FDR} < 0.05$; n.s.: $p_{FDR} > 0.05$.

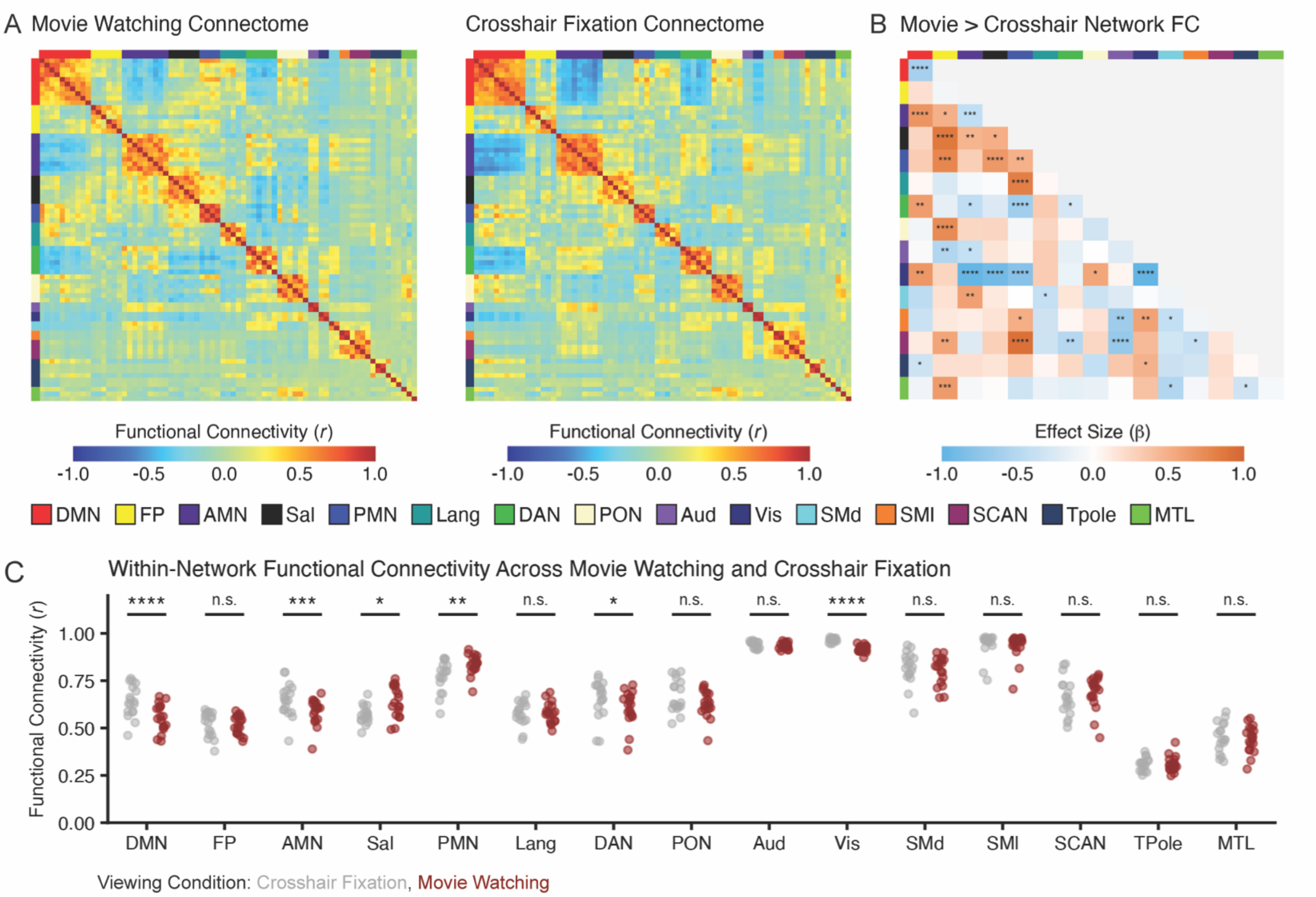


**Figure S14. Network functional connectivity between movie watching and crosshair fixation.** A) A movie watching and crosshair fixation connectome generated by parcellating the group average dense time series across MINT precision participants using the Masonic Institute for the Developing Brain Probabilistic Atlas (https://midbatlas.io/) (Hermosillo et al., 2024). B) The effect of viewing condition was quantified for each pattern of within- and between-network functional connectivity using linear mixed effects models with network functional connectivity (averaged across parcels assigned to network) as the outcome predicted by viewing condition (movies > crosshair) while covarying for fixed effects of age, sex, and head motion (mean framewise displacement in mm) and random subject intercepts. Probability values were false discovery rate corrected across the 120 models considered (each unique within- and between-network functional connectivity pattern). C) Scatterbar plots visualizing within-network functional connectivity values by viewing condition, significance of the viewing condition effect was quantified using the same linear mixed effects models following false discovery rate correction. *: $p_{FDR} < 0.05$, **: $p_{FDR} < 0.01$, ***: $p_{FDR} < 0.001$, ****: $p_{FDR} < 0.0001$.

**Table S1. Mapping Individual Networks Transdiagnostically (MINT) Precision Data, demographic and clinical characteristics of randomized participants by group and total.**

| Characteristic | Participant summary (*n* = 19 total) |
|---|---|
| *Age (Mean + SD [range])* | |
| Years | 15.8 ± 1.8 [12.7 to 17.9] |
| *Sex Assigned at Birth (n, % of total sample)* | |
| Female | 13 (68.42%) |
| Male | 6 (31.58%) |
| *Gender (n, % of total sample)* | |
| Girl | 12 (63.16%) |
| Boy | 6 (31.58%) |
| Nonbinary | 1 (5.26%) |
| *Ethnicity (n, % of total sample)* | |
| Hispanic or Latinx | 0 (0%) |
| Not Hispanic or Latinx | 19 (100%) |
| *Race (n, % of total sample)* | |
| Multiracial | 2 (11%) |
| White | 17 (89%) |
| *MINI Diagnosis (n, % of total sample)** | |
| Major Depressive Disorder | |
| Current | 8 (42%) |
| Past | 9 (47%) |
| Suicidality (current and lifetime) | 6 (32%) |
| Anxiety Disorders | |
| Panic | 1 (5%) |
| Separation Anxiety | 1 (5%) |
| Social Phobia | 2 (11%) |
| Specific Phobia | 2 (11%) |
| Generalized Anxiety | 8 (42%) |
| Post-Traumatic Stress Disorder | 1 (5%) |
| Obsessive Compulsive Disorder | 5 (26%) |
| Attention Deficit Hyperactivity Disorder (any type) | 9 (47%) |
| Tic Disorder | 6 (32%) |
| Oppositional Defiant Disorder (past 6 months) | 1 (5%) |
| Autism Spectrum Disorder (by history) | 9 (47%) |
| *Psychiatric Medications (n, % of total sample)* | |
| Selective serotonin reuptake inhibitors | 9 (47.37%) |
| Stimulants | 5 (26.32%) |
| Alpha-2 adrenergic agonists | 2 (10.53%) |
| Atypical antipsychotics | 2 (10.53%) |
| Other | 4 (21.05%) |
| *Child Behavior Checklist T-Score (Mean ± SD [Range])* | |
| *Broadband Scales* | |
| Internalizing | 62.0 ± 11.5 [34-80] |
| Externalizing | 46.6 ± 10.7 [34-69] |
| Total | 57.4 ± 9.3 [40-73] |
| *Syndrome Scales* | |
| Anxious Depressed | 63.5 ± 11.1 [50-86] |
| Withdrawn Depressed | 63.1 ± 9.7 [50-85] |
| Somatic Complaints | 60.7 ± 8.2 [50-80] |
| Social Problems | 56.4 ± 7.5 [50-72] |
| Thought Problems | 62.1 ± 7.6 [51-73] |
| Attention Problems | 58.5 ± 5.8 [50-66] |
| Rule Breaking Behavior | 52.5 ± 4.6 [50-64] |
| Aggressive Behavior | 53.5 ± 6.0 [50-70] |

* Diagnosis proportions (% of total sample does) does not sum to 100, as MINT precision participants could have multiple co-occurring conditions.

**Table S2. Healthy Brain Network Consortia Data, demographic and clinical characteristics of randomized participants by group and total**

| Characteristic | Participant summary (*n*=466 total) |
|---|---|
| *Age (Mean + SD [range])* | |
| Years | 12.8 + 3.5 [6.0 to 21.8] |
| *Sex Assigned at Birth (n, % of total sample)* | |
| Male | 248 (53%) |
| Female | 204 (44%) |
| Missing Data | 14 (3%) |
| *Ethnicity (n, % of total sample)* | |
| Not Hispanic or Latinx | 288 (62%) |
| Hispanic or Latinx | 84 (18%) |
| Unknown | 3 (1%) |
| Declined to answer | 15 (3%) |
| Missing data | 7 (16%) |
| *Race (n, % of total sample)* | |
| White/Caucasian | 208 (45%) |
| Black/African American | 49 (11%) |
| Hispanic | 36 (8%) |
| Asian | 12 (3%) |
| Indian | 2 (0%) |
| Native American | 0 (0%) |
| Alaskan Native | 0 (0%) |
| Native Hawaiian/ Other Pacific Islander | 0 (0%) |
| Two or more races | 67 (14%) |
| Other race | 9 (2%) |
| Unknown | 2 (0%) |
| Declined to answer | 2 (0%) |
| Missing data | 79 (17%) |
| *Scanning Site (n, % of total sample)* | |
| Rutgers University Brain Imaging Center | 175 (38%) |
| CitiGroup Cornell Brain Imaging Center | 291 (62%) |
| *Diagnosis (n, % of total sample*)* | |
| No diagnosis given | 54 (16%) |
| Major Depressive Disorder | 62 (19%) |
| Anxiety | 191 (58%) |
| Post-Traumatic Stress Disorder | 7 (2%) |
| Obsessive Compulsive Disorder | 19 (6%) |
| Attention Deficit Hyperactivity Disorder | 277 (84%) |
| Any Eating Disorder | 5 (2%) |
| Oppositional Defiant Disorder | 37 (11%) |
| Conduct Disorder Childhood Onset Type | 2 (1%) |
| Autism Spectrum Disorder | 76 (23%) |
| Missing data | 47 (10%) |
| *Child Behavior Checklist T-Score (Mean ± SD [Range])* | |
| *Broadband Scales* | |
| Internalizing | 57.3 ± 11.6 [33-88] |
| Externalizing | 53.4 ± 11.5 [33-87] |
| Total | 56.9 ± 10.9 [24-82] |
| *Syndrome Scales* | |
| Anxious Depressed | 58.6 ± 9.1 [50-92] |
| Withdrawn Depressed | 59.6 ± 9.8 [50-100] |
| Somatic Complaints | 57.7 ± 8.0 [50-90] |
| Social Problems | 57.0 ± 7.9 [50-88] |
| Thought Problems | 58.1 ± 8.1 [50-84] |
| Attention Problems | 61.4 ± 9.6 [50-97] |
| Rule Breaking Behavior | 55.5 ± 6.7 [50-91] |
| Aggressive Behavior | 57.5 ± 8.9 [50-88] |

* Diagnosis proportions (% of total sample does) does not sum to 100, as HBN consortia participants may have clinical consensus for up to 10 co-occurring diagnoses.

**Table S3. Data retention and quality between movie watching and crosshair fixation**

| Variable | Crosshair Fixation<br>Mean ± SD [range] | Movie Watching<br>Mean ± SD [range] | Effect size<br>*Cohen's d* [95% CI], $p_{1\text{-tailed}}$ |
|---|---|---|---|
| *MINT Precision Data (n=28 participants, prior to data quality exclusions)* | | | |
| Mean Framewise Displacement (mm) | 0.24 ± 0.33 [0.04 to 1.50] | 0.15 ± 0.12 [0.04 to 0.46] | 0.35 [0.03 to Inf], *p* = 0.04 |
| Retained Data at FD < 0.2 (minutes) | 30.28 ± 10.58 [0.79 to 39.48] | 31.85 ± 9.42 [2.55 to 39.56] | -0.28 [-Inf to 0.04], *p* = 0.08 |
| *HBN Consortia Data (n=1278 participants, prior to data quality exclusions)* | | | |
| Mean Framewise Displacement (mm) | 0.45 ± 0.60 [0.03 to 4.23] | 0.36 ± 0.41 [0.04 to 3.44] | 0.19 [0.14 to Inf], *p* < 0.001 |
| Retained data at FD < 0.2 (Minutes) | 6.37 ± 2.90 [0.13 to 10.00] | 6.39 ± 2.69 [0.13 to 10.00] | 0.01 [-0.04 to Inf], *p* = 0.37 |

**Table S4. Cortical network surface area estimates between movie watching and crosshair fixation, MINT precision data.**

| Network | Crosshair % of Total Cortical Surface<br>*M* ± SD [range] | Movie % of Total Cortical Surface<br>*M* ± SD [range] | Movie Clips > Crosshair Fixation<br>$\beta$ [95% CI], $p_{\text{2-tailed (FDR corrected)}}$ |
|---|---|---|---|
| Default Mode | 14.10 ± 2.16 [11.13 to 18.58] | 8.23 ± 1.71 [4.86 to 12.03] | -1.67 [-1.98 to -1.36], $p_{FDR} < 0.001$ |
| Action Mode | 12.22 ± 2.41 [8.47 to 17.11] | 9.00 ± 1.85 [5.87 to 12.14] | -1.21 [-1.58 to -0.84], $p_{FDR} < 0.001$ |
| Frontoparietal | 8.78 ± 1.48 [6.14 to 11.04] | 6.63 ± 1.59 [4.00 to 9.88] | -1.13 [-1.52 to -0.73], $p_{FDR} < 0.001$ |
| Visual | 12.50 ± 2.11 [7.34 to 15.92] | 10.87 ± 1.27 [8.42 to 13.83] | -0.81 [-1.39 to -0.23], $p_{FDR} = 0.010$ |
| Auditory | 4.01 ± 0.91 [2.66 to 5.71] | 3.08 ± 0.52 [2.51 to 4.70] | -1.06 [-1.50 to -0.62], $p_{FDR} < 0.001$ |
| Language | 4.99 ± 1.64 [1.39 to 8.67] | 9.82 ± 1.83 [6.12 to 12.38] | 1.61 [1.26 to 1.96], $p_{FDR} < 0.001$ |
| Parietal Memory | 3.97 ± 1.23 [2.27 to 7.58] | 6.49 ± 1.43 [3.81 to 9.79] | 1.37 [0.95 to 1.80], $p_{FDR} < 0.001$ |
| Salience | 4.40 ± 1.21 [1.74 to 7.30] | 7.44 ± 2.27 [4.12 to 11.88] | 1.24 [0.75 to 1.73], $p_{FDR} < 0.001$ |
| Parietal Occipital | 5.80 ± 1.62 [3.35 to 9.94] | 7.95 ± 1.48 [4.99 to 11.38] | 1.12 [0.59 to 1.65], $p_{FDR} < 0.001$ |
| Dorsal Attention | 9.68 ± 2.29 [5.41 to 13.97] | 12.30 ± 1.85 [7.81 to 16.78] | 1.04 [0.45 to 1.63], $p_{FDR} = 0.002$ |
| Somatomotor Lateral | 6.92 ± 1.75 [4.39 to 11.10] | 6.31 ± 1.46 [4.02 to 9.89] | -0.36 [-0.87 to 0.14], $p_{FDR} = 0.168$ |
| Somatomotor Dorsal | 4.86 ± 1.68 [2.45 to 8.12] | 4.06 ± 1.34 [2.30 to 6.54] | -0.49 [-1.00 to 0.02], $p_{FDR} = 0.081$ |
| Somato-Cognitive Action | 3.11 ± 1.15 [1.67 to 6.88] | 3.77 ± 0.82 [2.49 to 5.93] | 0.65 [0.27 to 1.02], $p_{FDR} = 0.004$ |
| Temporal Pole | 3.18 ± 1.54 [0.65 to 6.84] | 2.54 ± 1.03 [0.94 to 4.53] | -0.44 [-0.89 to 0.02], $p_{FDR} = 0.076$ |
| Medial Temporal | 1.48 ± 0.76 [0.64 to 3.41] | 1.52 ± 0.81 [0.47 to 3.39] | 0.08 [-0.30 to 0.46], $p_{FDR} = 0.667$ |

Network cortical surface area by viewing condition in the Mapping Individual Networks Transdiagnostically (MINT) precision data. Network cortical surface areas, % of the total cortical surface ($mm^2$), were derived using crosshair fixation and movie clips for each participant by viewing condition (28–38 minutes per condition, per participant). Network cortical surface areas are presented as mean ± standard deviation [range]. Effect sizes were calculated using linear mixed effects models with network cortical surface area as the outcome predicted by viewing condition (movie clips > crosshair fixation) while covarying for fixed effects of sex, motion (minutes retained), and age and random subject intercepts. Significance values were false discovery rate (FDR) corrected across the 15 mixed effects models. Effect sizes are reported as standardized beta coefficient [95% CI], probability value (FDR-corrected).

**Table S5. Cortical network surface area estimates between movie watching and crosshair fixation, HBN consortia data.**

| Network | Crosshair % of Total Cortical Surface<br>*M* ± SD [range] | Movie % of Total Cortical Surface<br>*M* ± SD [range] | Movie > Crosshair Effect Size<br>*β* [95% CI], $p_{2\text{-tailed (FDR corrected)}}$ |
|---|---|---|---|
| Default Mode | 11.80 ± 3.96 [1.25 to 23.03] | 7.77 ± 2.86 [0.09 to 16.12] | -0.95 [-1.06 to -0.85], $p < 0.001$ |
| Action Mode | 9.24 ± 3.55 [1.13 to 21.40] | 6.76 ± 2.89 [0.77 to 16.67] | -0.69 [-0.81 to -0.58], $p < 0.001$ |
| Frontoparietal | 8.10 ± 3.10 [0.34 to 18.85] | 9.12 ± 3.06 [2.13 to 20.04] | 0.37 [ 0.25 to 0.49 ], $p < 0.001$ |
| Visual | 10.11 ± 3.02 [0.48 to 18.62] | 5.97 ± 2.63 [0.81 to 15.58] | -1.15 [-1.25 to -1.04], $p < 0.001$ |
| Auditory | 3.67 ± 2.03 [0.33 to 12.94]] | 3.42 ± 1.51 [0.46 to 10.54] | -0.14 [-0.27 to -0.02], $p = 0.025$ |
| Language | 6.14 ± 3.09 [0.43 to 18.85] | 11.05 ± 2.89 [2.35 to 20.24] | 1.27 [1.18 to 1.37], $p < 0.001$ |
| Parietal Memory | 6.23 ± 2.74 [1.03 to 25.49] | 6.88 ± 2.82 [1.33 to 18.79] | 0.98 [0.88 to 1.09], $p < 0.001$ |
| Salience | 5.20 ± 2.51 [0.41 to 16.67] | 5.91 ± 2.57 [0.86 to 14.81] | 0.19 [0.08 to 0.31], $p = 0.002$ |
| Parietal Occipital | 7.56 ± 2.87 [1.20 to 19.49] | 8.57 ± 2.60 [1.36 to 18.28] | 0.28 [0.15 to 0.41], $p < 0.001$ |
| Dorsal Attention | 9.76 ± 3.51 [1.48 to 21.05] | 13.69 ± 3.68 [3.15 to 25.16] | 0.39 [0.26 to 0.51], $p < 0.001$ |
| Somatomotor Lateral | 8.57 ± 2.60 [1.36 to 18.28] | 9.05 ± 2.63 [3.91 to 25.01] | -0.30 [-0.40 to -0.19], $p < 0.001$ |
| Somatomotor Dorsal | 5.94 ± 2.78 [0.64 to 16.87] | 5.37 ± 2.31 [0.50 to 12.45] | -0.28 [-0.39 to -0.16], $p < 0.001$ |
| Somato-Cognitive Action | 2.95 ± 2.18 [0.02 to 15.44] | 2.99 ± 2.02 [0.06 to 14.56] | 0.02 [-0.09 to 0.13], $p = 0.731$ |
| Temporal Pole | 2.76 ± 2.09 [0.10 to 13.13] | 2.33 ± 1.76 [0.14 to 12.72] | -0.27 [-0.38 to -0.16], $p < 0.001$ |
| Medial Temporal | 1.00 ± 0.63 [0.14 to 7.75] | 1.12 ± 0.72 [0.15 to 6.09] | 0.08 [-0.03 to 0.18], $p = 0.154$ |

Network cortical surface area by viewing condition in the Healthy Brain Network (HBN) consortia data. Network cortical surface areas, % of the total cortical surface ($mm^2$), were derived using crosshair fixation and movie clips for each participant by viewing condition (8.01–9.99 minutes of post-censored data per participant, per condition). Network cortical surface areas are presented as mean ± standard deviation [range]. Effect sizes were calculated using linear mixed effects models with network cortical surface area as the outcome predicted by viewing condition (movie clips > crosshair fixation) while covarying for fixed effects of sex, motion (minutes retained), and age and random subject intercepts. Significance values were false discovery rate (FDR) corrected across the 15 mixed effects models. Effect sizes are reported as standardized coefficient [95% CI], probability value (FDR-corrected).
